# The desmosomal cadherin Desmogon is necessary for the structural integrity of the Medaka notochord

**DOI:** 10.1101/620070

**Authors:** Ali Seleit, Karen Gross, Michaela Woelk, Camilla Autorino, Jasmin Onistschenko, Lazaro Centanin

## Abstract

The notochord is an embryonic tissue that acts as a precursor to the spine. It is composed of outer sheath cells and inner vacuolated cells. Together they ensure the ability of the notochord to act as a hydrostatic skeleton until ossification begins. To date, there is still a paucity in our understanding of how the notochord cell types are specified and the molecular players controlling both their formation and maintenance remain poorly understood. Here we report that *desmogon*, a desmosomal cadherin, is essential for proper vacuolated cell shape and therefore correct notochord morphology. We trace *desmogon+* precursors and uncover an early developmental heterogeneity that dictates the balance of vacuolated and sheath cell formation. We demonstrate that the growth of vacuolated cells occurs asynchronously and reveal the presence of distinct injury sensing mechanisms in the notochord. Additionally, using a small-scale F0 CRISPR screen we implicate uncharacterized genes in notochordal integrity.

## Introduction

The notochord is the defining characteristic that unites all chordates (Lim et al. 2017; Satoh et al. 2012; Stemple 2005; Stemple et al. 1996; Corallo et al. 2018). Developmentally, it derives from the dorsal organizer region in vertebrate embryos (Stemple 2005; Ellis et al. 2013; Corallo et al. 2018). Subsequently, its constituent cells adopt a mesodermal fate and undergo convergent-extension movements (Satoh et al. 2012; Tada & Heisenberg 2012; Corallo et al. 2018; Stemple 2005; Stemple 2004). This results in a tube-like structure that runs along the anterior-posterior axis while simultaneously delimiting the dorso-ventral axis (Stemple 2005). In addition to providing structural support to embryos by acting as the major skeletal element during embryonic development (Stemple et al. 1996; Stemple 2005), it also has important signaling roles (Yamada et al. 1991; Yamada et al. 1993; Pourquie et al. 1993; Hebrok et al. 1998; Fouquet et al. 1997; Corallo et al. 2018; Stemple et al. 1996; Stemple 2005; Satoh et al. 2012). Indeed, patterning of adjacent tissues by the notochord is essential for correct morphogenesis to occur (Yamada et al. 1991; Yamada et al. 1993; Pourquie et al. 1993; Hebrok et al. 1998; Fouquet et al. 1997; Talbot et al. 1995; Stemple et al. 1996; Fleming 2004; Corallo et al. 2018). In vertebrates the notochord is a transient structure and a precursor to spine formation, it is eventually almost entirely replaced by vertebrae (Fleming 2004; Stemple 2005; Corallo et al. 2018; Lopez-Baez et al. 2018; Gray et al. 2014). Recently, it has been shown that correct spine patterning relies on segmentation cues present in the embryonic notochord (Wopat et al. 2018; Lleras Forero et al. 2018). This and other observations has led to the understanding that correct notochord morphogenesis is essential for normal spine formation (Fleming 2004; Lim et al. 2017; Wopat et al. 2018; Lleras Forero et al. 2018; Gray et al. 2014).

The notochord of all vertebrates studied to date (Ellis et al. 2014; Corallo et al. 2018) is formed of two cell-types: an outer sheath epidermal-like cell population and an inner vacuolated cell core (Stemple 2005; Corallo et al. 2018). The outer sheath cells cover the notochord tube and secrete ECM components that help in building and maintaining the peri-notochordal membrane (Ellis et al. 2013; Corallo et al. 2018; Yamamoto et al. 2010; Lim et al. 2017), while the inner-cells have large lysosomally-derived vacuoles that can withstand high hydrostatic pressure (Ellis et al. 2013; Ellis et al. 2014). This ability to act as a hydrostatic skeleton is particularly important in teleost fish as the strength and flexibility of the notochord is essential for proper locomotion – which in turn is necessary for survival – as embryogenesis concludes (Stemple et al. 1996; Ellis et al. 2014; Jiang & Smith 2007). A number of mutants affecting overall notochord formation and differentiation have previously been described in ENU screens (Stemple et al. 1996; Talbot et al. 1995). More recently, new players have been uncovered that are important for vacuolated cell formation and maintenance (Lim et al. 2017; Garcia et al. 2017). However, many decades after the first description of the notochordal cell types, there is still a paucity in our understanding of the molecular players that control the correct formation and maintenance of vacuolated and sheath cells and by extension the structural integrity of the notochord (Ellis et al. 2013).

Here we uncover new regulators of correct notochord morphogenesis in vertebrates. Desmogleins are a conserved family of calcium-binding cadherin transmembrane proteins. They localize to cellular membranes and are important for establishing strong cell-cell contacts and maintaining tissue integrity (Garrod & Chidgey 2008a). Structurally, they are part of the intercellular desmosome junctions (Garrod & Chidgey 2008a) and are expressed in tissues that undergo significant mechanical strain (Delva et al. 2009). We characterize the role of a fish-specific Desmoglein, hereafter referred to as *desmogon*, that is expressed in the notochord of Medaka and is to our knowledge the first Desmoglein reported to be present and functional in chordate notochords. Indeed, the loss of *desmogon* causes vacuolated cell defects and leads to structural deformities in the notochord. We generated a Tg(*desmogon*:EGFP) to address the origin and growth dynamics of vacuolated cells by 4D-imaging, a process revealed to be incremental and locally uncoordinated; vacuolated cells behave like autonomous units. Interestingly, *desmogon* also labels early disc-shaped notochord precursors that constitute a bipotent population. When analysed in 4D at the single-cell level, however, we reveal that each progenitor is unipotent and generates either one vacuolated cell or a number of sheath cells. In addition, exploiting the stable labelling of vacuolated cells in our Tg(*desmogon*:EGFP), we uncover two distinct types of regeneration responses in Medaka notochords, one spatially localized and the other global, that depend on the type of injury sustained. Finally, we use Tg(*desmogon*:EGFP) to carry out a small scale reverse-genetics screen on highly conserved and uncharacterized genes enriched in the notochord (Briggs et al. 2018; Farrell et al. 2018). Using this fast and straight-forward methodology we were able to implicate new players in correct notochord morphology in a vertebrate model. This work and approach, we believe, could lead to a deepening of our understanding of the origin of spine and vertebral defects.

## Results

### *desmogon* is a fish-specific desmosomal cadherin expressed in the notochord

While searching for a stable marker for neuromast border cells (Seleit et al. 2017) we serendipitously came across a novel uncharacterized *desmog-2-like* gene (ENSORLG00000017110), which we named *desmogon*. The 5.3Kb long transcript of *desmogon* is distributed over 14 exons and encodes a protein with at least 3 desmosomal cadherin domains and one cytoplasmic cadherin domain (Supplementary Figure 1). Based on the amino acid sequence the expected sub-cellular localization is plasma membrane and it is predicted to function as a component of the inter-cellular desmosome junctions. A list of all known orthologues of *desmogon* suggests that this gene is fish-specific, as it is absent in all other sequenced chordates (materials and methods for details) (Supplementary Table 1 and 2). Among fish, the *desmogon* locus is conserved in the vast majority of teleost branches although interestingly, it seems to have been lost in Zebrafish and Tetraodon (as evidenced by the syntenic conservation of the surrounding genomic region) (Supplementary Figure 1). *In situ* hybridization showed *desmogon* to be highly expressed in the developing notochord of Medaka (Figure 1A). To gain a better understanding of the dynamic spatial expression of *desmogon* we generated the Tg(*desmog*:EGFP) (Figure 1B-F) by using a 2.2kb proximal promoter region that contained strong peaks of H3K4 methylation (Supplementary Figure 1). Confocal analysis of mosaic, injected *desmogon*:EGFP and of Tg(*desmogon*:EGFP) medaka embryos revealed EGFP expression in the developing notochord throughout embryogenesis (Figure 1B-F, Supplementary Movies 1 & 2), this expression persists in adult fish in a segmented pattern along the spine (data not shown). Within the notochord, *desmogon* labels vacuolated cells (Figure 1 C, F, arrows in Figure 1E, 1F) and a proportion of covering sheath cells (yellow asterisks in Figure 1E, 1F). The expression of a Desmoglein family member in vacuolated and sheath cells suggests the presence of desmosomes in both cell-types, therefore we followed an electron microscopy (EM) approach to characterise the notochord of 10 dpf *wild-type* medaka larvae at sub-cellular resolution. Previous studies reported the existence of caveolae in the cellular membrane of vacuolated cells in the zebrafish notochord (Nixon et al., 2007; Lim et. al, 2017), which we confirmed is also present in medaka (Figure 1G, G’). Additionally, we observed the presence of desmosomes mediating the physical association of neighboring vacuolated cells (Figure 1G, G’), presumably to enhance their inter-cellular adhesion capacities. Desmosomes were also found connecting sheath to vacuolated cells (Figure 1H, I), and sheath to sheath cells (Figure 1I). Altogether, our results reveal the expression of an uncharacterised Desmoglein-like family member in the two cell types of the notochord that concurrently display desmosomes on their cellular membranes.

**Figure 1.**
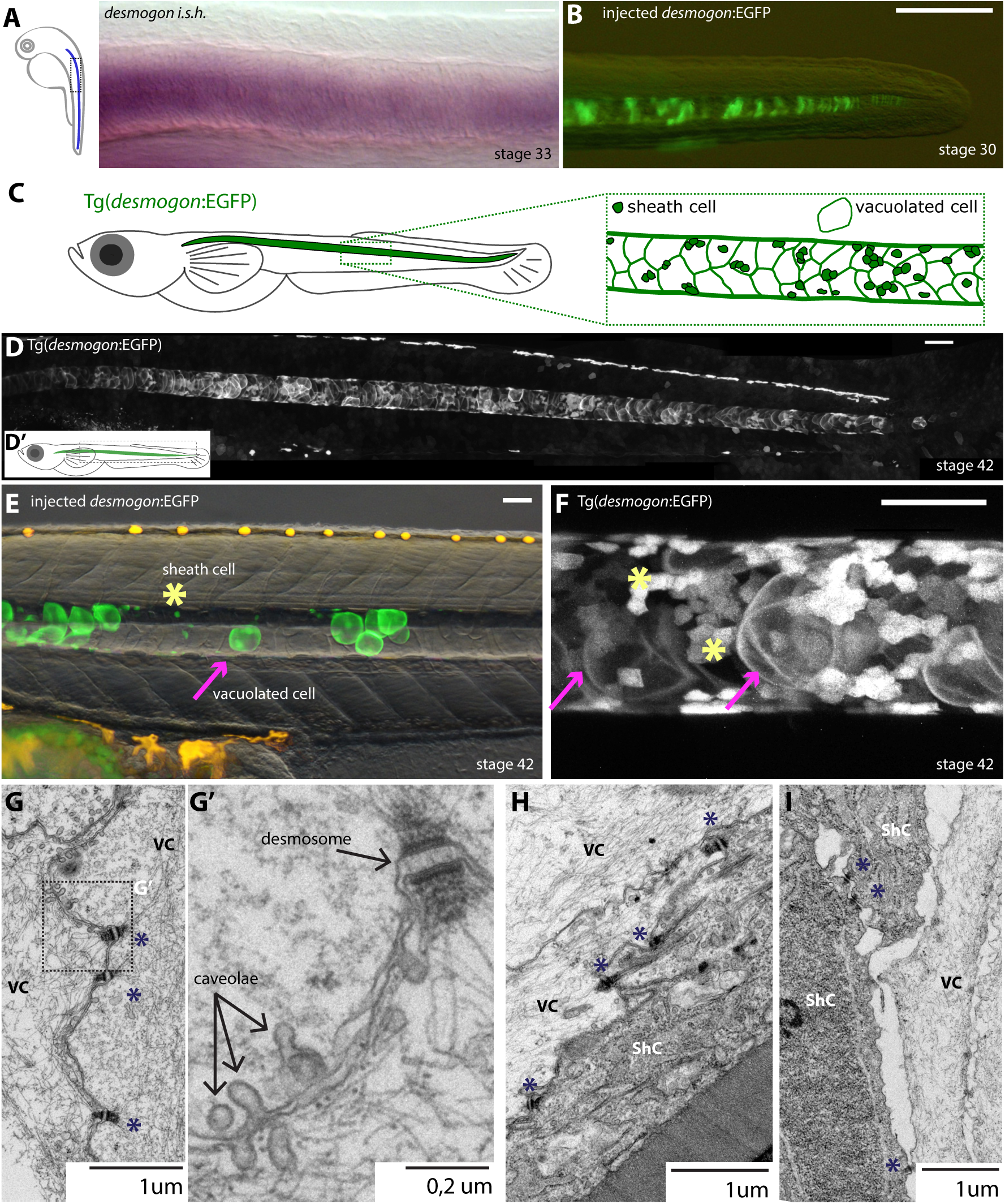
d*esmogon* is a desmosomal cadherin that labels the notochord throughout embryogenesis in Medaka. (A) *in-situ* hybridization on *desmogon* in stage 33 medaka embryos reveals strong enrichment in the notochord. Scalebar=20 microns (B) mosaic injection of desmogon:EGFP in medaka embryos stage 30, labels the notochord. Scale bar=100um (C-D) Transgenic line *Tg(desmog:EGFP)* labels the notochord in Medaka at stage 42 embryos. Scalebar=100 microns (E-F) Maximum projection of mosaic injected (E) and *Tg(desmog:EGFP)* (F) labelling notochord vacuolated cells and a proportion of covering sheath cells. Magenta arrows indicate vacuolated cells and yellow asterisks sheath cells. Scalebar= 50 microns (G) Longitudinal EM section between vacuolated cells connected by desmosomes (black asterisk) Scale bar= 1 micron (G’) Desmosomes and caveolae connecting vacuolated cells Scale bar = 0.2 micron. (H) Longitudinal EM section showing desmosomal connections between sheath cells and vacuolated cells scale bar= 1 micron. (I) Cross-section EM showing desmosomal connections between two sheath cells in addition to desmosomes between sheath cells and vacuolated cells scale bar= 1 micron.

### Extension of medaka notochord occurs via asynchronous vacuolated cell growth

The Tg(*desmog*:EGFP) allowed us to follow the axial extension of the notochord as the embryo develops. Notochord extension is mediated by the growth of vacuolated cells (Ellis et al., 2013; Corallo et al., 2018) that not only increase in size but also change their circular morphology into a more oblique shape as the notochord matures (Figure 2A, B) (N>10 notochords). During notochordal extension we observed that vacuole size in adjacent cells is not homogenous (Figure 2A), suggesting that the growth of vacuolated cells could be asynchronous. Indeed, by classifying vacuoles according to their size we were able to detect an intermingled distribution of vacuole area along the central, growing part of the notochord (Supplementary Figure 2A). We tracked vacuolated cell growth over time using transmitted, confocal and single plane illumination microscopy (SPIM) (Krzic et al., 2012) and observed that neighboring vacuolated cells grow anisotropically and asynchronously over time (Figure 2C, Supplementary Figure 2B for quantification) (Supplementary Movies 3, 4 and 5) in a process that appears to be irreversible (N>50 cells in 10 embryos). Interestingly, we also report that the global extension of the medaka notochord occurs in both the anterior and posterior directions. The growth of vacuolated cells in the central part of the notochord displaces anterior and posterior neighbors to their respective ends of the tube (Figure 2D-F’, Supplementary Movies 6-10). Overall, we show that the medaka notochord extends in a bidirectional manner, which is driven by a local asynchronous growth of vacuolated cells from the central section of the tube.

**Figure 2.**
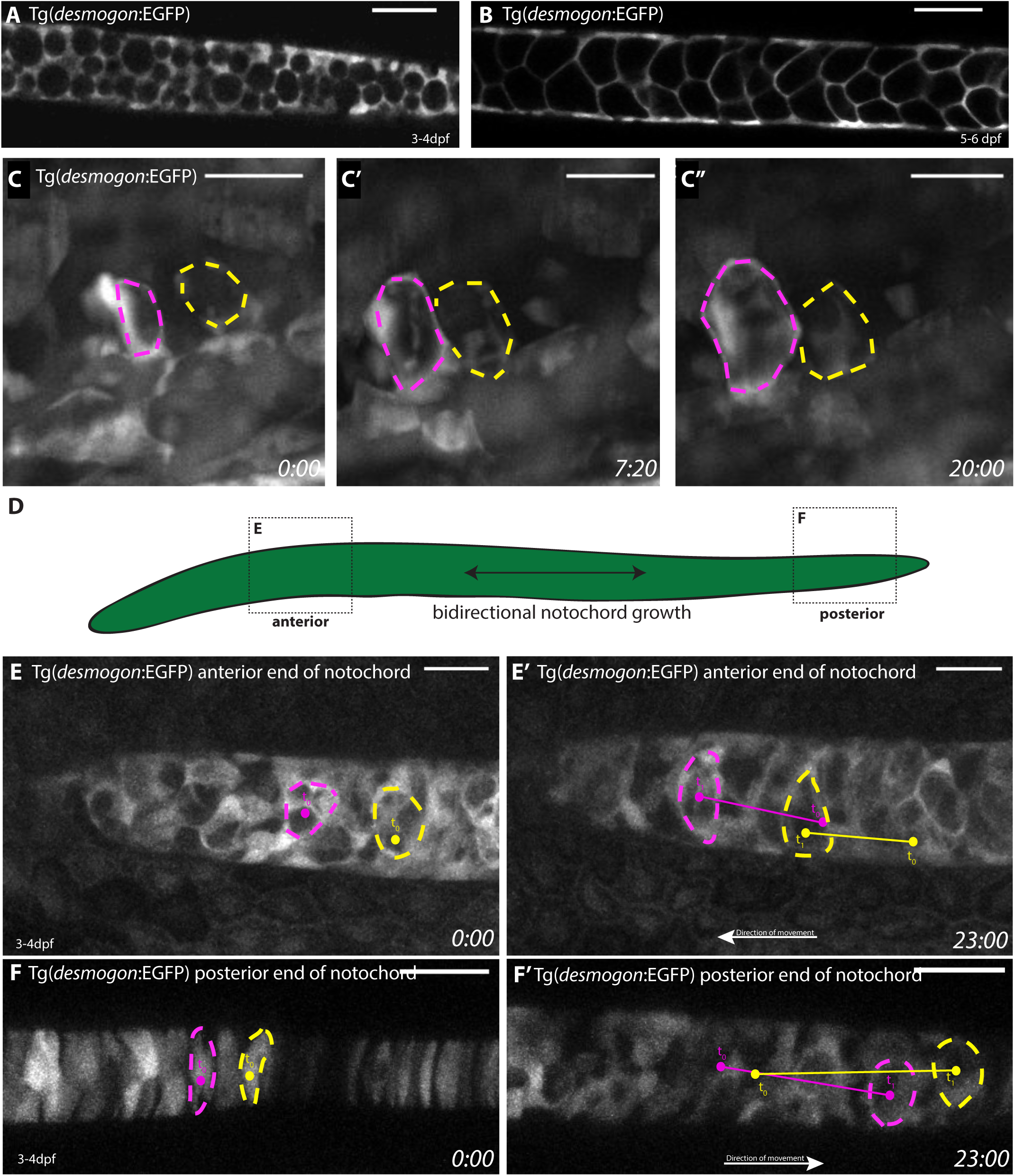
Live-imaging of *Tg(desmog:EGFP)* reveals bi-directional growth dynamics of the Medaka notochord. (A) Single plane of 3-4dpf *Tg(desmog:EGFP)* embryo showing vacuolated cells, notice more circular shape and small size compared to (B) 5-6dpf (*Tg)*desmogon:EGFP, bigger and oblique vacuolated cells. n>10. Scalebar=50 microns. (C-C’’) SPIM time-lapse imaging *of desmogon+* vacuolated cell growth over time highlighted with yellow and magenta asterisks (n>10 vacuolated cells in 3 embryos at 4-5dpf and n>10 vacuolated cells in 2 embryos at 3-4dpf). Scalebar=20microns. (D) Schematic diagram of different sectors of the notochord at the anterior and posterior tip, the growth of the tube is driven bi-directionally from the mid-section. (E-E’) Vacuolated cells at the anterior end of notochord are displaced more anteriorly as the tube grows from the mid-section. Two outlined vacuolated cells at the anterior end of the growing notochord tube are labelled at t0 (magenta and yellow dotted lines) and traced over time t1 distance between t0 and t1 indicative of displacement more anteriorly. Scalebar=30 microns. Time in hours. (F-F’) Disc-shaped precursors at the posterior end of notochord are displaced more posteriorly as the tube grows from the mid-section. Two disc-shaped precursors at the posterior end of the growing notochord tube are labelled at t0 (magenta and yellow dotted lines) and traced over time t1 distance between t0 and t1 indicative of displacement more posteriorly. Scalebar=30 microns. Time in hours. N=3 independent embryos.

### Unipotency of *Desmogon+* disc shaped precursors

The extension of the notochord is complemented by the addition of new differentiated cells at the posterior tip, originating from so-called disc-shaped precursors (Dale and Topczewski 2011; Yamamoto et al., 2010; Melby et al.,1996). The population of disc-shaped-precursors was previously shown to generate both differentiated cell types (Yamamoto et al., 2010; Melby et al., 1996), and therefore constitutes a bi-potent population. While analysing sparsely labelled notochords of medaka larvae that were injected with *desmogon*:EGFP (Figure 3A, B) at the two cell stage, we noticed that clusters tended to contain either vacuolated or sheath cells (N=41 sheath cells clusters, N=39 vacuolated cell clusters, N=80/93 cell-type specific cluster and N=13/93 clusters containing both cell types, N=93 clusters in 26 mosaic larvae). These results suggest that under physiological conditions disc-shaped precursors might be bi-potent as a population, but fate-restricted as individual cells. To test this hypothesis directly, we followed the process by 4D imaging. Disc precursors are labelled in the Tg(*desmogon*:EGFP) (Figure 1B, Figure 3 A and C, Supplementary Movie 10 and 11), which allowed us to follow the notochordal differentiation process dynamically at the single cell level. We observed two distinct, mutually exclusive cellular behaviours in *desmogon+* disc-shaped precursors. On the one hand, they can directly generate a single vacuolated cell (Figure 3C-C’’, magenta dot) (Supplementary Movie 10-12) (N=17/23 cells in 3 embryos). These cells did not divide throughout our imaging and therefore constitute a post-mitotic cell type (N=3 embryos at 4 dpf, and N=2 embryos at 3dpf image for 24h, N>50 cells) (Supplementary movie 3), as was reported for zebrafish vacuolated cells (Garcia et al., 2017). On the other hand, disc-shaped precursors can also undergo a dorso-ventral symmetric division leading to the exclusive formation of sheath cells (Figure 3C-C’’ yellow dots) (Supplementary Movie 12) (N=6/23 cells in 3 embryos), indicating that disc-shaped precursors are unipotent in medaka.

**Figure 3.**
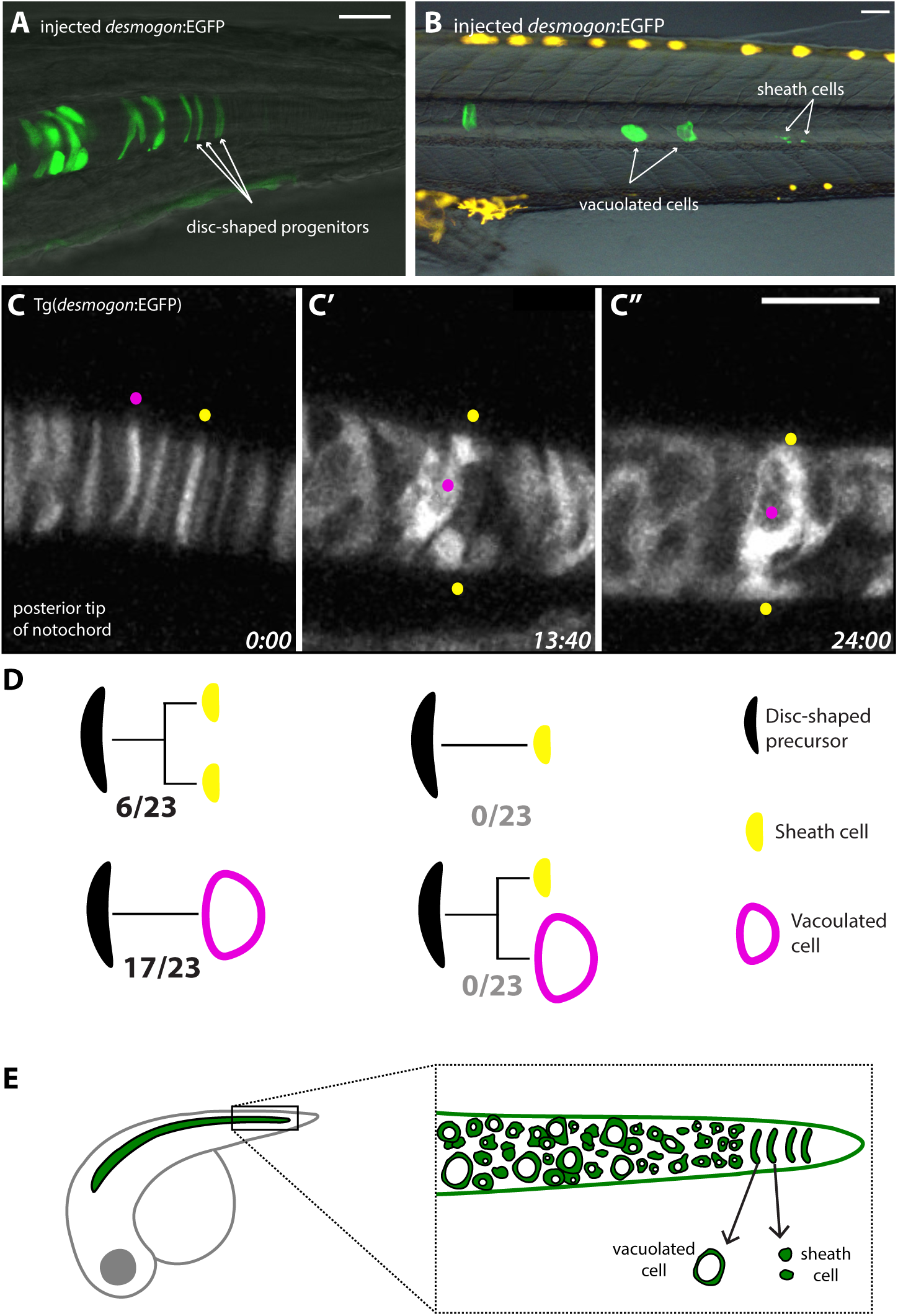
Bi-potent disc-shaped precursor population is fate-restricted at single cell level. (A) Mosaic injected medaka embryo *desmog:EGFP* showing the presence of disc-shaped precursors at the posterior tip of the growing notochord tube. (B) independent clones of sheath cells and vacuolated cells occur in mosaic injected fish. (C-C’’) Time-lapse imaging of posterior tip of growing Medaka notochord shows the differentiation dynamics of disc-shaped precursors. Two disc-shaped precursors are labelled (magenta and yellow dot). The magenta labelled precursors start differentiating into a vacuolated cell while the yellow dot labelled precursor undergoes a dorso-ventral division leading to the formation of two sheath cells. Time in hours (D) Disc-shaped precursors at the growing tip of the medaka notochord are fate-restricted and either generate vacuolated cells without dividing or sheath cells after dorso-ventral division. Total disc-shaped precursors followed= 23 in 3 embryos. Scale bars are 20um in A, 1mm in B, and 30 um in C.

To test whether the observed unipotency of disc-shaped precursors is a medaka-specific feature we decided to explore the same process in zebrafish. Even though a *desmogon* orthologue does seem to be present in *Danio rerio*, we observed EGFP-labelled disc-shaped precursors, vacuolated and sheath cells in *desmogon*:EGFP zebrafish injected embryos (Supplementary Figure 3, and Supplementary Movie 13). This indicates that the transcriptional machinery driving expression of medaka *desmogon* is conserved in distantly related teleost fish. Live-imaging of embryos with EGFP+ clones in the undifferentiated notochord revealed that similar dynamics govern the differentiation process in zebrafish as in Medaka. Disc-shaped *desmogon+* precursors directly differentiate into vacuolated cells (N=8 cells in 6 embryos) that will not undergo mitosis throughout our imaging (N>10 cells in 6 embryos). Additionally, a closer look at the notochordal differentiation dynamics in zebrafish revealed that the birth and growth of vacuolated cells is governed by the same dynamics as we report in Medaka (from disc-shaped precursors that differentiate and grow incrementally in a locally unsynchronized manner) (N>10 cells n=6 embryos) (Supplementary movie 13-15). Additionally, and as we report for medaka, disc-shaped precursors in zebrafish form sheath cells exclusively after undergoing a dorso-ventral symmetric division (N=16 cells in 6 embryos) (Supplementary Figure 3, Supplementary movie 14 and 15). In contrast to the post-mitotic nature of vacuolated cells, sheath cells continue to divide long after acquiring their characteristic morphology and peripheral position. Interestingly in both medaka and zebrafish we also observed the presence of newly formed sheath cells that do contain small vacuoles (n>10 cells in n=6 embryos in Zebrafish and n=3 embryos in Medaka) (Supplementary Movie 14 and 16), suggesting that this feature could reflect morphologically distinct sub-populations of sheath cells. Overall, our results show on the one hand that *desmogon*:EGFP is a suitable tool to study early aspects of notochord differentiation in distantly related teleosts, and demonstrates the presence of unipotent, fate-restricted disc-shaped precursors that exclusively generate either vacuolated or sheath cells. Since both vacuolated and sheath cells come from unipotent precursors and since we and others have shown that these disc-shaped precursors are exhausted by the end of notochordal development (Ellis et al. 2013; Corallo et al., 2018), we wondered whether and how vacuolated cells can be replaced after injury of mature Medaka notochords.

### Local regenerative response after targeted vacuolated cell loss in medaka

Recent findings using a number of injury paradigms have reported that zebrafish can efficiently regenerate the notochord (Garcia et al. 2017; Lopez-Baez et al., 2018). Given the differences in regenerative capacities among teleosts (Lust & Wittbrodt 2018; Ito et al. 2014; Lai et al. 2017), we used Tg(*desmogon*:EGFP) to address the response to local notochord injuries in medaka. Spatially targeted and precise multi-photon laser ablation of 6-10 vacuolated cells of 5-6 dpf Tg*(desmogon:*EGFP) embryos resulted in the specific loss of cells in the area of injury (Figure 4A-A’’, Supplementary Figure 4 Supplementary movie 17 and 18). Both sheath and vacuolated cells outside the ablated zone retain a normal morphology and the overall integrity of the notochord is unaffected (Figure 4A-B, entire Z-stacks in Supplementary Movie 19 and 20). Two days post injury we observed the appearance of small *desmogon+* vacuolated cells specifically in the area of injury (Supplementary Figure 4, Supplementary movie 19) (n=8 embryos). Interestingly, the overall morphology, presence of a small vacuole, and EGFP+ expression of these cells was highly reminiscent of the earliest vacuolated sheath cells we observed during development in both medaka and zebrafish notochords, suggesting that the regenerative response is mediated by the same cell type in both species (Garcia et al. 2017). The small vacuolated cells grow in size over time as assessed at 5 days post injury (Figure 4C, Supplementary movie 20) (N=8 embryos), this growth followed the same asynchronous rationale we observed under physiological conditions in Medaka. Overall, our results indicate that Medaka notochords can mount a robust and local regeneration response to vacuolated cell loss, which is spatially restricted to the initial injury site.

**Figure 4.**
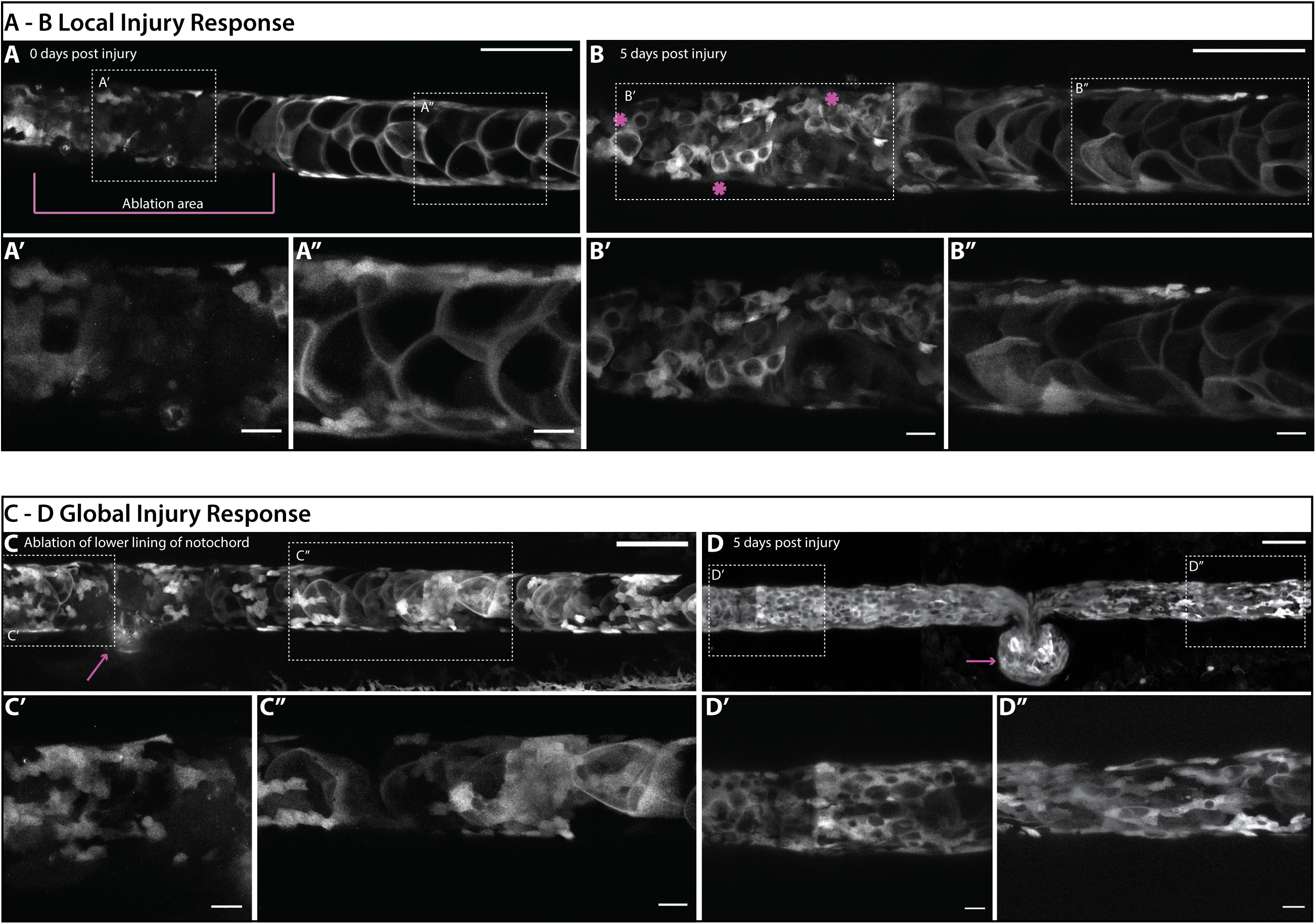
Local and global regeneration dynamics after notochord injury. (A-A’’)*Tg(desmog:EGFP)* 6dpf embryo directly post laser ablation of vacuolated cells. Ablation area indicated by magenta lines. Vacuolated cells in the ablated zone are missing. Vacuolated cells outside ablation zone are intact and morphologically normal. Scalebar= 100 microns. (B-B’’) Same embryo 5 days post injury, notice the presence of small vacuolated cells specifically in the site of injury, vacuolated cells grow in size as compared tp 48 hours post injury see supplement, notochord is intact. Scalebar= 100 microns. n=8 embryos. (C-C’’) Ablation of vacuolated cells and lower lining of notochord tube. (D-D’’) 5 days post injury magenta arrow highlights growing leakage of *desmogon+* vacuolated cells outside of the notochord. Failure to repair and correctly regenerate lower tube lining is evident. Gross morphological defects apparent over entire length of notochord. Notice the appearance of small vacuolated cells throughout the notochord. Scalebar= 100 microns. n=3 embryos.

### Global regeneration responses following peri-notochordal membrane injury

The perinotochordal membrane is a thick ECM layer that ensheathes the notochord and helps to maintain its integrity (Ellis et al. 2013; Corallo et al. 2018; Yamamoto et al. 2010; Lim et al. 2017). To test the effect of a sudden loss of hydrostatic pressure within the notochordal tube, we ablated the lower lining of the peri-notochordal membrane (Figure 4 C-C’, Supplementary Figure 4, Supplementary movie 21). Two days post-injury we observed *desmogon+* cells leaking outside of the notochord tube, indicating that the lower lining of the notochordal membrane failed to be repaired (Supplementary Figure 4). Given the post-mitotic nature of vacuolated cells and the dynamics of regeneration shown earlier, the local leakage at the initial injury site decreases the number of vacuolated cells that remain in the tube. This was accompanied by the appearance of small desmogon+ cells containing a vacuole as reported for the local response, although here these are found anterior and posterior to the initial targeted injury site (Figure 4 C-C’’, Supplementary Figure 4). Five days post-injury the leakage of notochord vacuolated cells continued, forming a herniated structure, and resulting in a significant perturbation of notochord morphology. Small vacuolated cells persisted anterior and posterior to the injury site (Figure 4D-D’’) (n=3). Overall, we conclude that injury to the peri-notochordal membrane, which cannot be efficiently repaired in Medaka, leads to cell leakage and a perturbed notochord morphology. This in turn triggers a global regeneration response that is not spatially restricted to the initial injury site.

#### Targeted CRISPR screen uncovers novel regulators of notochord integrity

In addition to structural damage sustained by the notochord due to targeted laser ablation, we wondered whether similar phenotypes can be observed by genetic perturbations. It has recently been reported that F0 CRISPR phenotypes concur with the ones observed in stable mutant lines (Wu et al. 2018; Lischik et al. 2018; Trubiroha et al. 2018). We therefore decided to use the Tg*(desmogon:*EGFP) as a fast and straight-forward read-out of notochordal defects. Exploiting the recently generated single-cell transcriptome data from Zebrafish (Briggs et al. 2018; Farrell et al. 2018) we chose a number of well annotated and poorly characterized genes that were strongly conserved across vertebrates and highly expressed during Zebrafish notochord morphogenesis (*arrdc3a, kcnk6, pmp22b, si:dkey-261h17.1 and vgll2b*) and designed 2 gRNAs targeting selected exons for each gene (for details of selection criteria see materials and methods). For *vgll2b* this resulted in 55% of injected embryos showing morphological defects in notochord shape and integrity including twisting and bending of the notochord tube (Figure 5B-B’’, Table 1). Targeting *arrdc3a* resulted in 30% of injected embryos showing disruption in notochordal integrity, with strong phenotypes including buckling and kinking (Figure 5C-C’’ and Table 1). Targeting *kcnk6, si:dkey-261h17.1, pmp22b* with the same approach resulted in strong phenotypes on the notochord during embryogenesis (Supplementary Figure 6) but additionally all three showed significant pleiotropic effects (Stemple et al. 1996), including general growth retardation, shorter body axes and gross morphological defects (for quantifications on all injections and the phenotypes observed see Table 1). In addition, we also targeted the teleost specific *desmogon* with 3 gRNAs to address whether it has any functional role during notochord morphogenesis and/or maintenance (Supplementary Figure 5). Phenotypes in *desmogon* crispants included the loss of notochord integrity and shape as revealed by kinking and buckling along the notochord tube (Figure 5D-E’, Table 1). In conclusion, using a straight-forward reverse-genetics approach we have uncovered novel and conserved regulators of notochord morphogenesis and maintenance in vertebrates. The use of the Tg(desmogon:EGFP) line allowed us to complement our gross description of the notochordal phenotypes with a more detailed view on the cellular organization of the tissue (Figure 5 F). We noticed that unlike the other candidate genes *desmogon* crispants specifically exhibited defects in vacuolated cell morphology (Figure 5 E’). We therefore decided to take a closer look at the cellular defects in *desmogon* mutant notochords.

| Gene Name | Full name | proposed roles structure and localization | CRISPR Injected F0 Phenotypes % | Phenotypic description |
| --- | --- | --- | --- | --- |
| *dsgon* | *desmogon* | constituent of desmosomal structures, plasma membrane | 40,6 | local collapses of vacuolated cells, notochord bending, local notochord collapses, no pleiotropic effects, highly specific to notochord, 90% of embryos survive to stage 42 |
| *vgll2b* | *vestigial-like family member 2b* | predicted transcription factor TF/co-factor, nuclear | 55 | strong phenotypes in the notochord, bending and twisted notochords, spirals of notochord, notochord tube coiling, local collapses of notochord no big pleiotropic effects, 79% of embryos survive to stage 42 |
| *arrdc3a* | *arrestin domain containing 3a* | regulating signal transduction at G protein-coupled receptors | 30 | wavy notochords, twisted notochords, no strong pleiotropic effects, 77% of embryos survive to stage 42 |
| *kcnk6* | *potassium two pore domain channel subfamily K member 6* | ion transport, plasma membrane | 44,3 | kinks and wavy notochords, shorter axis elongation, general growth retardation 73% survive to stage 42 |
| *si:dkey-261h17.1* | ENSORLG00000015828 | cell adhesion, membrane bound | 42,1 | notochord spirals and twists, wavy notochord, kinks, perturbed axis elongation, 69% of embryos survive to stage 42 |
| *pmp22b* | *peripheral myelin protein 22b* | helical transmembrane, integral component of membranes, | 31,2 | strong pleiotropic effects, growth retardation and developmental delays, heart edemas, smaller eyes, wavy notochords, spiral notochords and twists, short axis/failure of axis extension, patterning defects, 58% survive to stage 42 |

**Figure 5.**
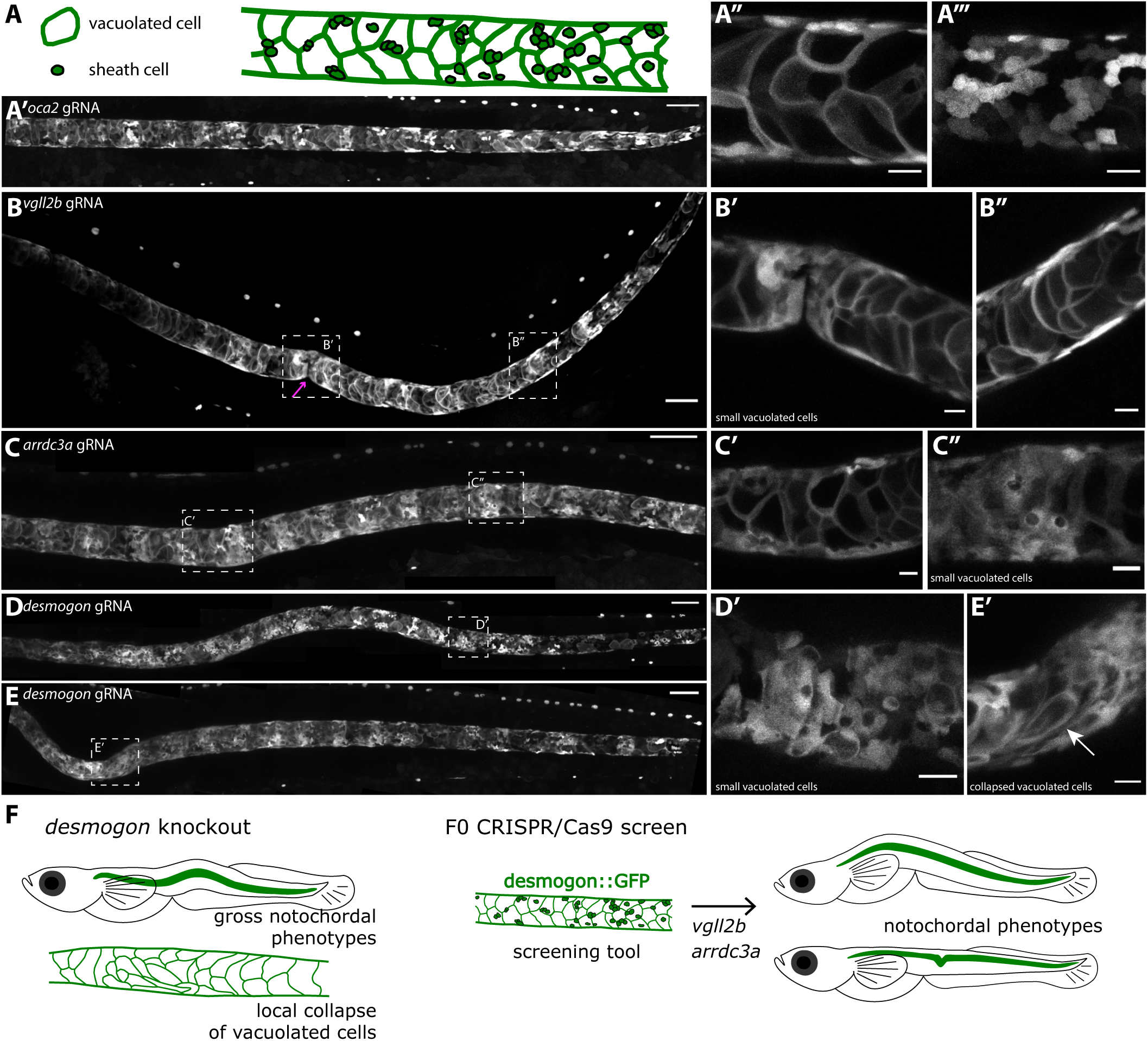
Gross morphological defects in the notochord of *vgll2b, arrdc3a and desmogon* CRISPR injected embryos. (A) Overall morphology of control *oca2* gRNA & Cas9 injected notochord in *Tg(desmogon:EGFP)*(B-B’’) *vgll12b* gRNA1,2 & Cas9 injected into *Tg(desmog:EGFP)* results in morphological defects in the notochord. Notice magenta arrow where notochord is twisted, overall notochord bending observed. Scale bar=100microns. Embryos with phenotypes in notochord 66/120. 79% of embryos survive to stage 42. (C-C’’) *arrdc3a* gRNA1,2 & Cas9 injected into *Tg(desmog:EGFP)* results in notochord bending. Scalebar=100microns. Embryos with phenotypes 29/97. 77% of embryos survive to stage 42. (D-F) Strong phenotypes in *desmogon* CRISPR injections with gross morphological defects in notochord integrity and notochord buckling and bending. maximum projections. Scalebar=100 microns. Embryos with notochord phenotypes 65/160. Over 90% of embryos survive to stage 42.

### *Desmogon* crispants and stable mutants exhibit notochordal lesions of collapsed vacuolated cells

CRISPR/Cas9 injection into Tg*(desmogon:*EGFP) resulted in lesions containing collapsed vacuolated cells along the length of the notochord, which were absent in embryos injected with control *oca2* gRNAs (Figure 6 A-C, Supplementary Movies 24, 25) (Lischik et al. 2018). The vast majority of injected embryos survived until stage 42 and no obvious pleiotropic effects were detected, suggesting a notochord-specific role for Desmogon. A proportion of F0 injected embryos that survived to adulthood showed strong signs of bending and defective spine formation (data not shown). We therefore decided to perform alizarin-red bone stainings on *desmogon* crispants and mutants. This revealed the presence of defects in vertebrae (smaller, misshaped, and fused vertebrae) (Figure 6 F-H), linking proper vacuolated cell shape conferred by Desmogon to correct spine formation in Medaka. The described phenotypes for the F0 injected *desmogon* crispants were consistently recapitulated in *desmogon* mutants (Figure 6A-D, Supplementary Movies 22, 23 and Supplementary Figure 5 for alleles isolated in the stable mutant line). A closer analysis of the phenotypes affecting embryos with collapsed notochords revealed the presence of larger lesions that were consistently filled with *desmogon*+ small vacuolated cells (Supplementary Movie 22 & 23). To gain a better understanding of the structural phenotypes observed in *desmogon* mutants, we decided to perform electron microscopy (EM) on mutant notochords. We first focused on the presence and structural integrity of mutant desmosomes, and could not observe any obvious phenotype when compared to *wild-type* desmosomes (Figure 7 A-C). Longitudinal sections on wild-type notochords were characterised by the typical highly ordered array of vacuolated cells (Figure 7D). This contrasts with the structural disorganization present in lesioned areas of the *desmogon* mutant notochords (Figure 7 E-E’). EM data also showed the presence of vacuolated cells of appreciably different sizes (Figure 7 E-E’, arrows), evidence of vacuolated cell collapse (Figure 7F-F’), and invasion of sheath cells into the central notochord tube that indicates the possible triggering of a regenerative response in the lesioned area (Figure 7 E-G asterisk). In addition to being a marker for vacuolated cells, we therefore believe that *desmogon* has a functional role in proper notochord integrity and shape. Its loss leads to vacuolated cell collapse and the appearance of lesions that contain small vacuolated cells, this can in turn lead to gross morphological defects in the notochord.

**Figure 6.**
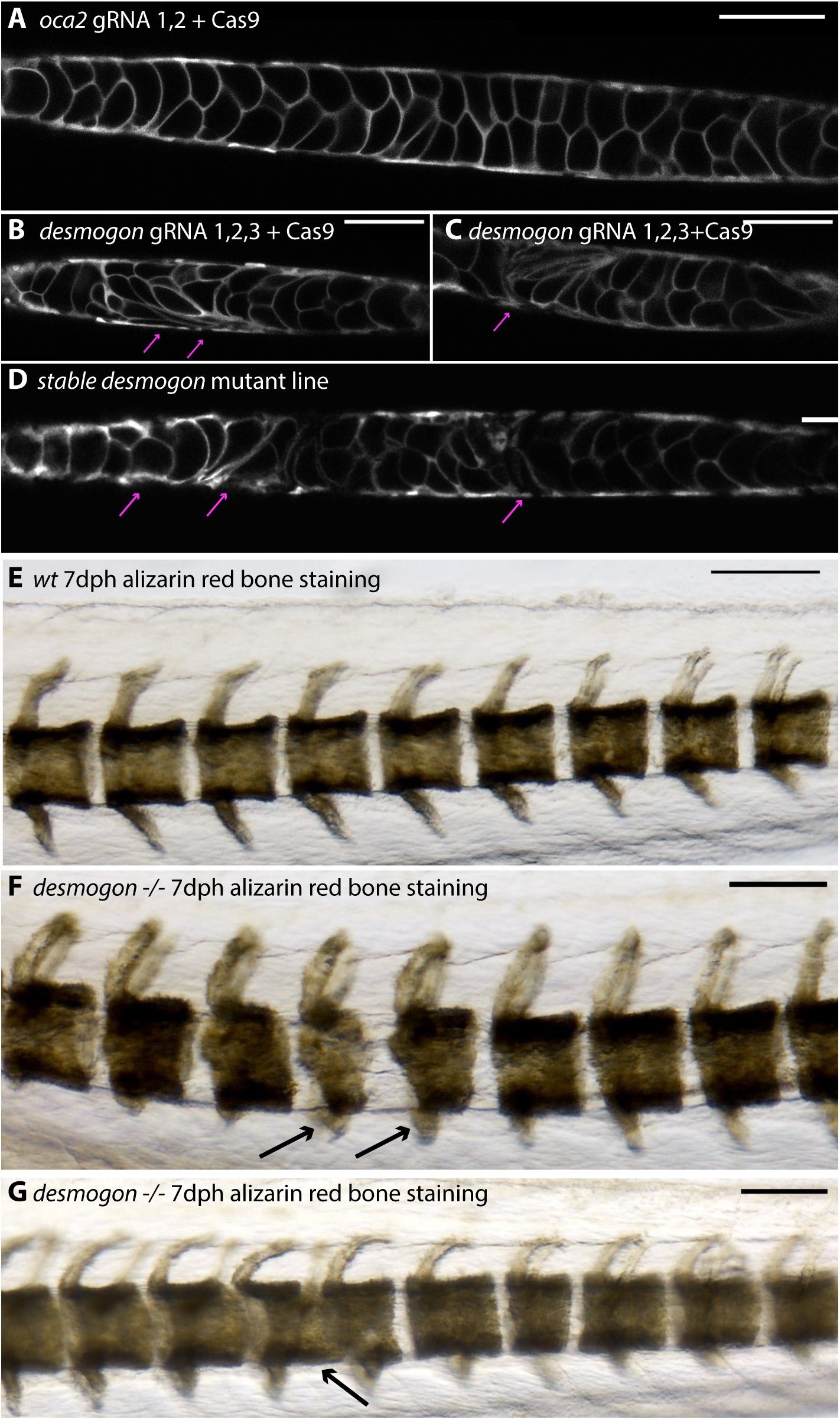
*desmogon* mutants exhibit notochordal lesions and vacuolated cell collapse. (A)Control CRISPR injected *Tg(desmog:EGFP)* with o*ca2* gRNA1,2 & Cas9. Single plane. Scalebar=100 microns. (B, C) *desmogon* gRNA1, 2, 3 & Cas9 injected into *Tg*Desmogon:EGFP results in local collapse of vacuolated cells and lesions in the notochord. Single plane. Scalebar=100 microns. (D) Stable *desmogon* CRISPR mutant line recapitulates phenotypes observed in the injected generation Scalebar=100microns. (E) Alizarin red bone staining on *wt* 7days post hatch Medakas shows the highly ordered and regularly sized and spaced vertebral segmentation. (F-G) Alizarin red bone staining on *desmogon* Crispants and stable mutants shows the presence of defects in vertebral size (black arrows in F) and fused vertebrae (black arrows in G). Scale bars on E, F and G= 30 microns.

**Figure 7.**
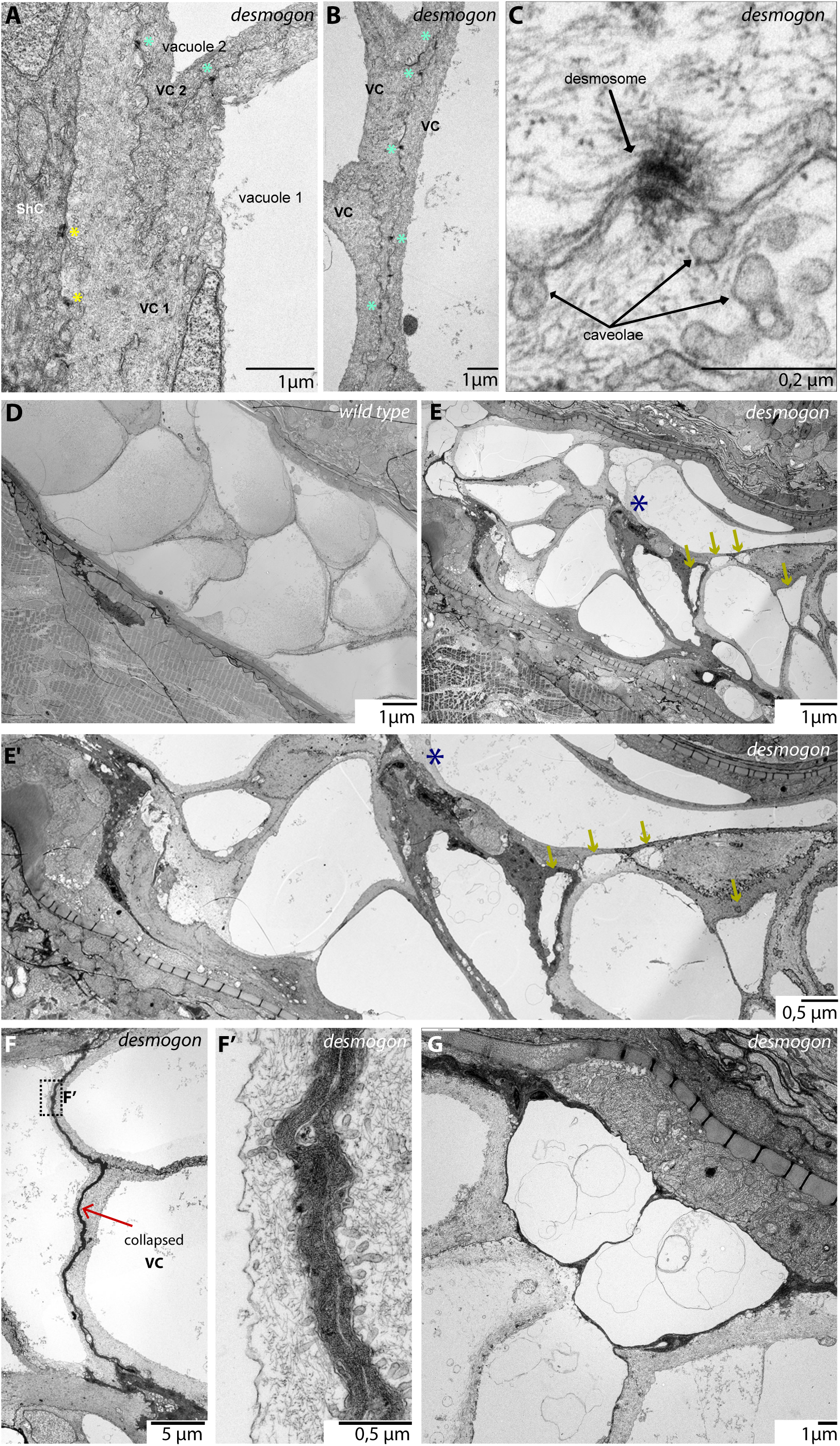
EM on *desmogon* mutants shows normal desmosomes but disrupted vacuolated cellular morphology and evidence of a regenerative response. (A-C) EM imaging reveals the presence of desmosomes between vacuolated and sheath cells, and vacuolated cells in *desmogon* mutants. Scale bar= 1 micron in A and B. Scale bar=0.2 microns in C. (D) Longitudinal EM section through wild-type stage 42 medaka notochords with highly ordered vacuolated cell arrangement (E) Longitudinal EM section through lesioned *desmogon* mutant stage 42 notochords, notice the structural disorganization, vacuolated cells with varying sizes (yellow arrows), invading sheath cells (black asterisk) and evidence of collapsed vacuolated cells. (F-F’) Longitudinal EM section on *desmogon* mutant notochords reveals the presence of collapsed vacuolated cells. (G) Small vacuolated cells are present in Longitudinal sections of EM in *desmogon* mutants and can be clearly distinguished from the neighboring vacuolated cells and sheath cell.

Interestingly, the position, number and size of lesioned areas in the notochord (both in the injected and stable mutant line) were variable among siblings. We therefore wondered whether the appearance and position of the vacuolated cell defects in *desmogon* mutants depended on the mechanical stress the notochord endures during tail movement and bending, as has been recently reported for caveolae mutants in Zebrafish (Lim et al., 2017). This could be particularly relevant in Medaka due to the long embryogenesis, larger overall size and more tightly confined space within a stronger chorion as compared to zebrafish. Importantly, Medaka embryos regularly bend their tails during development (Iwamatsu 2004). To test whether movement is required for the appearance and/or severity of the collapsed vacuolated cell phenotype, we injected *alpha-bungarotoxin* mRNA into *desmogon* mutants (Lischik et al. 2018). This led to the transient loss of movement throughout embryogenesis that was later restored at 9-10 dpf when the toxin levels dampened. Injected embryos and un-injected controls were removed from the chorion 3dpf and followed day by day for the appearance of notochordal lesions. In total, 21/21 alpha-bungarotoxin injected *desmogon* mutant did not show any signs of movement and did not display any collapsed vacuolated cell phenotypes throughout embryogenesis. This contrasted with the earlier appearance of notochordal lesions in un-injected *desmogon* mutants (with tail movements) on day 6-8 post fertilization. Of the 21 alpha-bungarotoxin injected *desmogon* mutants, 16 showed lesions of collapsed vacuolated cells in the notochord only upon regaining the tail movement, 2 fish died before any visible movement and did not show any lesions and 3 fish did not show any visible lesions even after movement. The results strongly suggest that the collapse of vacuolated cells occurs after the specification and growth phases and that the phenotype largely depends on, and is exacerbated by, the mechanical stress induced by tail bending and movement. This is in line with previous reports in other mutants affecting vacuolated cell shape and integrity in zebrafish (Lim et al., 2017). Overall, we report that Desmogon is a necessary desmoglein in maintaining proper vacuolated cell morphology specifically during the intense mechanical stress imposed by the physiological movement of Medaka fish.

## Discussion

Desmogleins are a conserved family of desmosomal cadherins that localize to the plasma membrane of cells (Garrod & Chidgey 2008b; Delva et al. 2009). They are a constituent part of desmosomes and are important mediators of strong inter-cellular adhesion. Indeed, desmosomes have been shown to be expressed in cell types that operate under significant mechanical strain (Garrod & Chidgey 2008b; Delva et al. 2009). Loss of desmosomal function leads to disruption of tissue integrity (Garrod & Chidgey 2008a). The notochord is a tissue that is constantly assailed by strong mechanical stresses (Lim et al. 2017; Garcia et al. 2017; Corallo et al. 2018). It is able to withstand those pressures primarily because of its structural organization: large vacuolated cells on the inside of the tube are surrounded by a strong peri-notochordal membrane formed by sheath cells (Lim et al. 2017; Yamamoto et al. 2010; Koehl et al. 2000; Adams et al. 1990). Our study on *desmogon*, a fish-specific desmosomal cadherin, is the first showing a desmoglein family member expressed and functional in vertebrate notochords.

### *desmogon* is necessary for correct notochord morphology in Medaka

The structural demands on teleost notochords are particularly high given that larvae need to swim and feed as soon as embryogenesis concludes (Jiang & Smith 2007; Stemple et al. 1996; Ellis et al. 2014). Significantly, this happens before the ossification and formation of spines (Corallo et al. 2018; Lleras Forero et al. 2018; Wopat et al. 2018; Fleming 2004; Gray et al. 2014). This could explain the specific allocation of this desmoglein family member expression to the notochord. To test whether *desmogon* has a functional role in the Medaka notochord we targeted it by CRISPR/Cas9 and observed flattened vacuolated cells and lesions along the length of the notochord. This could either be a sign of collapsed vacuolated cells (Lim et al., 2017) or a failure of vacuolated cells to properly form. Our EM data on lesioned mutant notochords and the *alpha-bungarotoxin* experiment strongly suggest that the phenotype results from the local collapse of vacuolated cells due to movement. Interestingly, in areas that contained larger lesions (both in injected fish and in stable mutants), we consistently observed the appearance of small vacuolated cells. This is highly reminiscent of results we report from the regeneration experiments and suggests that the collapse of *desmogon* mutant vacuolated cells can trigger a regenerative response. Indeed, EM results on lesioned notochords shows evidence of invading sheath cells. Our results indicate that *desmogon* is necessary for the maintenance, but not the formation or growth, of vacuolated cells. Paralyzed desmogon mutant embryos showed no signs of notochordal disruption or vacuolated cell defects. In general, we observed a strong correlation between the size of the mutant lesion and the structural integrity of the notochord, bigger lesions invariably led to buckling and kinking of the notochord tube. Our results are in line with previous observations that have shown that correct vacuolated cell morphology is essential for the notochord to withstand the high mechanical stresses it faces (Lim et al. 2017; Ellis et al. 2013; Garcia et al. 2017; Fleming 2004; Adams et al. 1990). Failure of properly building up and maintaining the high hydrostatic pressure inside the notochord tube leads to bending and buckling along the length of the notochord (Ellis et al. 2013; Corallo et al. 2018). Intriguingly, the cellular phenotype we report for *desmogon* mutants in Medaka is highly reminiscent of the caveolae mutant phenotype in zebrafish (Lim et al. 2017). This suggests that the notochord as a system might have multiple independent buffering capacities against vacuolated cell collapse under mechanical stress. Given the essential role of the notochord in the early survival of teleost larvae it is possible that these buffering capacities evolved to imbue the notochord with more strength by acting synergistically and buffering against the malfunctioning of one component. Lastly, we show that lesioned vacuolated cells in *desmogon* mutant notochords lead to a disruption of proper vertebral segmentation complementing recent findings in the field (Lim et al. 2017; Wopat et al. 2018; Lleras Forero et al. 2018; Gray et al. 2014). Our results demonstrate that the presence of *desmogon* in the Medaka notochord is necessary for correct vacuolated cell shape and by extension proper notochord (and spine) morphology and integrity.

The fact that *desmogon* is present in the vast majority of teleost branches argues that it has a conserved role in notochord maintenance in fish. However, there are two intriguing exceptions, *desmogon* has no detectable orthologues in Tetraodon or in Zebrafish. It seems likely that both fish have lost *desmogon* through the course of their evolution, this interpretation is supported by the fact that we can identify syntenic genomic regions that do not contain *desmogon.* It remains a formal possibility, however, that an orthologue with a highly divergent nucleotide sequence exists in Zebrafish and Tetraodon. It would be interesting to see whether other desmosomal cadherins have taken over the notochordal role of *desmogon* in those species. Indeed, other desmosomal cadherin family members have been implicated in notochord integrity in Zebrafish (Goonesinghe et al. 2012), although mutants display earlier gastrulation defects that complicate the proper characterisation of notochordal phenotypes. Interestingly, injecting our medaka *desmogon* partial promoter driving GFP in Zebrafish results in vacuolated and sheath cell labelled clones. This suggests that the core transcriptional machinery driving tissue-specific expression of *desmogon* remains in place and active in Zebrafish. It would be of interest to investigate when the deployment of desmosomal cadherins in notochords arose during evolution and how wide-spread its usage is among the different chordate clades.

### *Tg(desmogon:EGFP)*as a screening tool for genes involved in proper notochord morphology

It has been reported that most biomedical research today focuses heavily on a limited number of genes, leaving behind potentially important genes understudied (Stoeger et al. 2018). Making use of our newly generated transgenic line in combination with the recently published and publicly available single cell transcriptomics data from vertebrate embryos (Briggs et al. 2018; Farrell et al. 2018), we attempted to address this imbalance. To do so we performed a small scale F0 CRISPR screen, the efficacy of which has been recently demonstrated in Zebrafish (Wu et al. 2018) and confirmed in other fish species (own observations and personal communications). Indeed, recent work in Medaka (Lischik et al. 2018), in addition to our results from *desmogon* mutants, argues for the use of F0 injected embryos as a method to analyse tissue-specific phenotypes. Briefly, we focused on conserved well-annotated genes that were highly and differentially expressed in developing notochords. We addressed whether they might be involved in proper notochord morphology by using the *desmogon:GFP* line as a fast and straight-forward read-out for notochord shape and integrity. Broadly, the targeted genes fell into two categories. *kcnk6, si:dkey-261h17.1, pmp22b* constituted the first category and showed strong pleiotropic effects in addition to notochordal defects. Delineating the cause of the pleiotropy is difficult given the essential signaling role of the notochord. (Yamada et al. 1991; Yamada et al. 1993; Pourquie et al. 1993; Hebrok et al. 1998; Fouquet et al. 1997; Corallo et al. 2018; Stemple et al. 1996). The observed defects could either arise from the inability of the notochord to correctly pattern adjacent tissue or from notochord independent roles for these genes during embryogenesis. This makes assigning causal phenotypes more difficult. The second group of targeted genes contained the conserved putative co-transcriptional factor *vgll2b* and the highly conserved membrane bound arrestin *arrdc3a.* Both genes showed specific defects in notochord morphology and structure when targeted and no overt pleiotropic phenotypes. This argues for functional roles for these genes that are likely to be notochord specific. The precise cellular defects caused by these two genes remain unclear although in both cases vacuolated cells appeared morphologically normal. It is clear, though, that targeted notochords appear unable to withstand the high mechanical strain and buckle under pressure. It remains a challenge for the future to decipher the genetic networks of these genes and to assess whether their roles are functionally conserved in higher vertebrates. All in all, using a simple, targeted, reverse-genetics approach, in combination with publicly available data from single cell transcriptomics (Briggs et al. 2018; Farrell et al. 2018), we believe we have implicated new players in correct vertebrate notochord integrity. This methodology can be easily adapted to other contexts and promises to aid in the study of neglected genes with potentially important functions.

### Notochord vacuolated cells during development

We made use of the newly generated *Tg(desmogon:EGFP)* line to address the formation and growth dynamics of notochord vacuolated cells during development. It has previously been reported that one vacuole exists per vacuolated cell under homeostatic conditions (Ellis et al. 2014) and therefore vacuole growth can be used as a proxy for cellular volume growth. Our results confirm that vacuolated cells grow in volume over time as has been reported before (Ellis et al. 2013; Ellis et al. 2014) and reveal that they do so anisotropically, changing their morphology in the process from more roundish to more oblique shapes. This might be due to the increased cellular packing as the notochord expands. While it has been previously shown that notochords extend over time and that this supports axis elongation in vertebrate embryos (Ellis et al. 2014; Garcia et al. 2017; Ellis et al. 2013), it is still unclear exactly how this growth and expansion is coordinated. A possible mechanism could be that a morphogen gradient synchronizes the growth of neighbouring cells in an orderly fashion. Our dynamic data, in both medaka and zebrafish embryos, argues against the presence of such a signal. We reveal that the growth of vacuolated cells in Medaka and Zebrafish is an incremental one-way process that is locally uncoordinated; neighbouring vacuolated cells grow at different rates. This strongly suggests cell-autonomous mechanisms of vacuolated cell growth, how this is coordinated globally to eventually reach an equivalent size remains unclear and constitutes an interesting avenue for future research. It could be that the final size reached is close to the physiological limit as has been previously suggested (Ellis et al. 2014). On the global growth of the notochordal tissue we report a previously unrecognized bi-directional growth mode where the growth initially driven in the mid-section displaces the posterior and anterior segments of the tube to their respective ends.

In addition to incrementing their size cell-autonomously we report that the number of vacuolated cells increases as the notochord grows. However, in line with previous observations from Zebrafish (Garcia et al. 2017), we have observed no cell division of vacuolated cells. To characterize the initial steps of vacuolated cell formation during notochord development we employed a highly temporally resolved 4-D approach. We observed that vacuolated cells arise from disc-shaped precursors as previously reported (Melby et al. 1996; Dale & Topczewski 2011). Sparse labelling of these *desmogon*+ precursors in developing Zebrafish notochords combined with long-term live-imaging revealed a hitherto unrecognized *in vivo* behavioural heterogeneity of the disc-shaped precursors. Either they directly differentiate into vacuolated cells (forgoing any division), or they generate sheath cells by undergoing a dorso-ventral symmetric division, that could be followed by further amplifying rounds of mitosis. When considering the position along the AP axis, the developmental time and the levels of EGFP+ expression, we were unable to reliably predict the output of a precursor cell. It seems plausible therefore that the decision to form a vacuolated or sheath cell is not predetermined nor controlled by tissue-level morphogens, but rather depends on sensing the needs of the growing notochord tube in a local manner. Indeed, it has been previously reported that Notch-Jag1 signalling is important in the balance between vacuolated and sheath cell formation (Yamamoto et al. 2010), suggesting that fate acquisition could be resolved locally among neighbouring cells. Complementarily, it is possible that local mechanical forces that arise during the expansion of the tube could operate on inherently plastic disc-shaped precursors and contribute to adopting vacuolated or sheath cell identity.

### Notochord vacuolated cells in regeneration

It has been recently shown that Medaka hearts and retinas have a vastly reduced regenerative potential as compared to Zebrafish (Lust & Wittbrodt 2018; Ito et al. 2014; Lai et al. 2017). However, we have previously reported that Medaka neuromasts are able to regenerate efficiently (Seleit, et al. 2017), and the same happens after mechanical amputations on the caudal fin (Katogi et al. 2004) and injuries to the liver (Van Wettere et al. 2013). It therefore seems that Medaka has a highly variable tissue-specific regenerative capacity (Kang et al. 2016). We wondered whether Medaka can regenerate lost notochord vacuolated cells. It has been shown in Zebrafish that loss of *caveolae* mutant vacuolated cells in response to mechanical strain triggers an efficient regeneration response (Garcia et al., 2017). This is mediated by sheath cells that invade the inside of the notochord tube and trans-differentiate into vacuolated cells (Garcia et al., 2017). Localized laser-ablation of vacuolated cells in Medaka leads to a very similar process as in Zebrafish where small vacuolated cells specifically invade the injury site. Interestingly, these invading cells start growing in size asynchronously mirroring the un-coordinated nature of growth that we observe for vacuolated cells during development. The appearance of small vacuolated cells during regeneration is another reminiscent feature of what we report during notochord development. Indeed, a proportion of sheath cells formed from dividing precursors during development do initially contain small vacuoles that highly resemble the cells participating in the regenerative response. This suggests two things; a) the presence of sub-populations of sheath cells that display distinct behaviours both during development and regeneration (Lopez-Baez et al., 2018), b) sheath cells participating in the regenerative response most likely re-acquire a small vacuole and could thus be reverting to an earlier state in their developmental history. Reactivation of developmental programs has been a hallmark of efficient regeneration in a variety of models (Tanaka & Galliot 2009; Rodrigo Albors et al. 2015; Kaloulis et al. 2004) and it seems likely that this takes place in the notochord of Medaka after vacuolated cell loss.

The highly specific spatially localized response to vacuolated cell injury we observe in Medaka indicates that there are mechanisms in place that can sense injured tissue without the need to activate a global regeneration program (LoCascio et al. 2017). Indeed, it has been shown that the release of vacuolated cell contents upon apoptosis can trigger a local regenerative reaction from neighbouring sheath cells (Garcia et al. 2017). A similar process can therefore be occurring in Medaka. Interestingly, when we ablated a small part of the peri-notochordal membrane this led to vacuolated cell leakage at the injury site. As the vacuolated cell leakage continued we witnessed the activation of a global regenerative response anterior and posterior to the original injury site. This could be observed by the presence of small vacuolated cells along the entire length of the notochord. It therefore seems likely that sheath cells can also respond to injury without specific vacuolated cell death, suggesting that sheath cells might be able to sense other stresses like tissue tension. It has recently been reported in Zebrafish that a *wilms+* subpopulation of sheath cells gets activated in response to a coarse needle injury to the notochord (Lopez-Baez et al. 2018). This sub-population mainly forms scar-tissue that acts as a stopper to maintain notochordal integrity (Lopez-Baez et al. 2018). In Medaka, leakage of vacuolated cells at the site of injury indicates that the peri-notochordal membrane was not efficiently repaired. It is tempting to speculate that this is due to the absence or delayed activation of the wilms+ subpopulation of sheath cells. Overall, our results strongly argue for the presence of distinct injury-sensing mechanisms in sheath cells (dependent and independent of vacuolated cell death). We also report the existence of two regenerative responses in Medaka notochords, one spatially localized and the other global, that depend on the type of injury sustained. Our data positions vacuole re-acquisition by sheath cells as the key step for replenishing vacuolated cells regardless of the type of injury sustained. Identifying the molecular trajectories sheath cells traverse will allow a better understanding of the existing heterogeneities among and plasticity of sheath cells both during development and regeneration. This will in turn allow a more targeted exploitation of the potential of sheath cells in treating notochordal and by extension spinal cord defects in vertebrates.

## Legends of Supplementary Files

### Supplementary Movies

**Supplementary movie1** *Tg(desmog:EGFP)* labels the Medaka notochord. *Tg(desmog:EGFP)* stage 42 Medaka embryo. Z-stack shows labelled vacuolated cells throughout the notochord and a proportion of sheath cells labelled. Scalebar=100 microns.

**Supplementary movie2** 3D reconstruction of *Tg(desmog:EGFP*) Medaka notochord. stage 42 Medaka embryo. 3D projection shows the specific labelling of the notochord tube by 2.2kb *desmogon* promoter region. Scalebar=100 microns.

**Supplementary movie3 and 4** Asynchronous growth of neighboring vacuolated cells. *Tg(desmog:EGFP)* 4-5dpf embryo. Time-lapse SPIM imaging of growing *desmogon+* vacuolated cells (3). Time-lapse confocal imaging of growing *desmogon+* vacuolated cells (4). Notice the anisotropic and autonomous nature of vacuolated cell growth, n>10 vacuolated cells in 3 embryos at 4-5dpf and n>10 vacuolated cells in 2 embryos at 3-4dpf. Scalebar= 20microns. Time in hours.

**Supplementary movie5** Bright-field imaging of developing mid-section of medaka notochord shows asynchronous growth of vacuolated cells. 5dpf embryo. Scale bar 30 microns. Time in hours.

**Supplementary movie6** Bi-directional growth of the notochord. Anterior section of the developing notochord extends anteriorly. *Tg(desmog:EGFP)* 4-5dpf embryo. Time-lapse imaging of developing anterior section of the notochord reveals a push towards the anterior end. No observable divisions of *desmogon+* vacuolated cells. n=3 embryos at 4-5dpf and n=2 embryos at 3-4dpf. Scalebar=20 microns. Time in hours.

**Supplementary movie7** Bi-directional growth of the notochord. Bright-field imaging of anterior section of the notochord reveals push towards the anterior end. Scale bar= 30 microns. Time in hours.

**Supplementary movie8** Bi-directional growth of the notochord. Bright-field imaging of posterior tip of developing notochord shows that the tube extends posteriorly. Scale bar= 30 microns. Time in hours.

**Supplementary movie9** Bi-directional growth of the notochord. Anterior section of the developing notochord shows individual cells displaced more anteriorly in *Tg(desmog:EGFP)* 5dpf embryo. Scale bar= 20 microns. Time in hours.

**Supplementary movie10** Disc-shaped precursors are located at the posterior tip of the developing Medaka notochord and are pushed posteriorly by the growing tube. *Tg(desmog:EGFP)* 4-5dpf embryo near extending posterior end of the tail. *desmogon+* disc-shaped precursors can differentiate into vacuolated cells that grow in size over time. Time in hours. Scalebar=30 microns.

**Supplementary movie11** 3-D reconstruction of disc-shaped precursors. Tg(*desmog:EGFP)* injected *3*dpf embryo near extending posterior end of the tail. Notice the labelling of undifferentiated disc-shaped precursors.

**Supplementary movie12** Unipotency of disc-shaped precursors in developing medaka notochords. *Tg(desmog:EGFP)* 4-5dpf embryo near extending posterior end of the tail. *desmogon+* disc-shaped precursors can either differentiate into vacuolated cells that grow in size over time or undergo a dorso-ventral division to produce two sheath cells. Time in hours. Scalebar=30 microns.

**Supplementary movie13** *desmog:EGFP* labels vacuolated and sheath cells in developing zebrafish notochords. Clones of *desmogon+* cells in 24hpf Zebrafish embryo developing notochord. Both sheath cell clones and vacuolated cell clones are present. Notice the growth of vacuolated cells is locally uncoordinated in Zebrafish as is the case in Medaka. Scalebar=30 microns. Time in hours.

**Supplementary movie14** Unipotency of disc-shaped precursors in zebrafish. Clones of *desmogon+* disc shaped precursors in 24hpf Zebrafish embryo developing notochord. Direct differentiation of the anterior disc-shaped precursor into a vacuolated cell. Posterior disc shaped precursor produces two sheath cells after a dorso-ventral division Scalebar=30 microns. Time in hours.

**Supplementary movie15** Dorso-ventral division of disc shaped precursors leads to sheath cell production. Clones of *desmogon+* disc shaped precursors in 24hpf Zebrafish embryo. Notice the presence of small vacuoles in newly formed sheath cells, that disappear over time. Scalebar=30 microns. Time in hours.

**Supplementary movie16** Presence of small vacuoles in newly formed sheath cells in developing Medaka notochords. Tg(*desmog:EGFP) 4-5*dpf embryo near the mid section of the notochord. Notice the presence of multiple sheath cells with small vacuoles some of which undergo mitotic divisions. Scalebar=20 microns. Time in hours.

**Supplementary movie17** Pre-injury Z-stack through notochord. *Tg(desmog:EGFP)* 6dpf embryo. Pre-Ablation stack through notochord. Scalebar= 100 microns.

**Supplementary movie18** Post-injury Z-stack through notochord shows precise ablation of vacuolated cells. *Tg(desmog:EGFP)* 6dpf embryo. Post-laser ablation stack through notochord. Notice the loss of vacuolated cells anteriorly at the site of injury. Scalebar= 100 microns.

**Supplementary movie19** 2 days post-injury Z-stack through notochord reveals presence of small vacuolated cells specifically at the site of injury. *Tg(desmog:EGFP)* embryo. 48hours post-injury stack through notochord. Small vacuolated cells appear specifically in the site of injury. Scalebar= 100 microns.

**Supplementary movie20** 5 days post-injury Z-stack through notochord reveals the growth of small vacuolated cells specifically at site of injury. *Tg(desmog:EGFP)* embryo. 5 days post-injury stack through notochord. Vacuolated cells grow in size specifically in the site of injury. Scalebar= 100 microns.

**Supplementary movie21** Injury to peri-notochordal membrane. *Tg(desmog:EGFP)* embryo. Post-ablation stack through notochord. A number of vacuolated cells and the lower lining of the notochord ablated. Scalebar= 100 microns.

**Supplementary movie22** Lesions of collapsed vacuolated cell in desmogon Crispants. *Tg(desmog:EGFP)* 6-7dpf desmogon gRNA1,2,3 & Cas9 injection. Z-stack through notochord. Strong phenotypes show abnormal notochord lesions. Notice the presence of small vacuolated cells and cellular debris. Overall a disorganized notochord structure is evident. Scalebar= 50 microns.

**Supplementary movie23** Lesions of collapsed vacuolated cells in *desmogon* stable mutants. *Tg(desmog:EGFP)* 6-7dpf desmogon F1 mutant. Z-stack through notochord. Strong phenotypes show abnormal notochord lesions. Phenotype resembles what is observed in the injected generation. Scalebar= 30 microns.

**Supplementary movie24** Control z-stack through *oca2* gRNA injected Tg(desmog:EGFP) embryos. 6-7dpf *oca2* gRNA1,2+ Cas9 injection. Z-stack through notochord. Notice normal notochord and vacuolated cell morphology. Scalebar= 100 microns.

**Supplementary movie25** *desmogon* gRNA injected embryos exhibit lesions of collapsed vacuolated cells. *Tg(desmog:EGFP)* 6-7dpf Desmogon gRNA1,2,3 & Cas9 injection. Z-stack through notochord. Local collapses of vacuolated cells are evident and vacuolated cell morphology is highly perturbed. Scalebar= 100 microns.

### Supplementary Table

**Table 1** List of genes targeted by CRISPR and quantification of phenotypes in F0 injections. For injection numbers *vgll2b* 66/120 injected fish showed the described phenotype for *arrdc3a* 29/97 for *desmogon* 65/160 for *pmp22b* 50/160 for *si:dkey261h17.1* 16/38. Representative phenotypes are shown in supplementary figures.

### Supplementary Figures

**Supplementary Figure 1.**
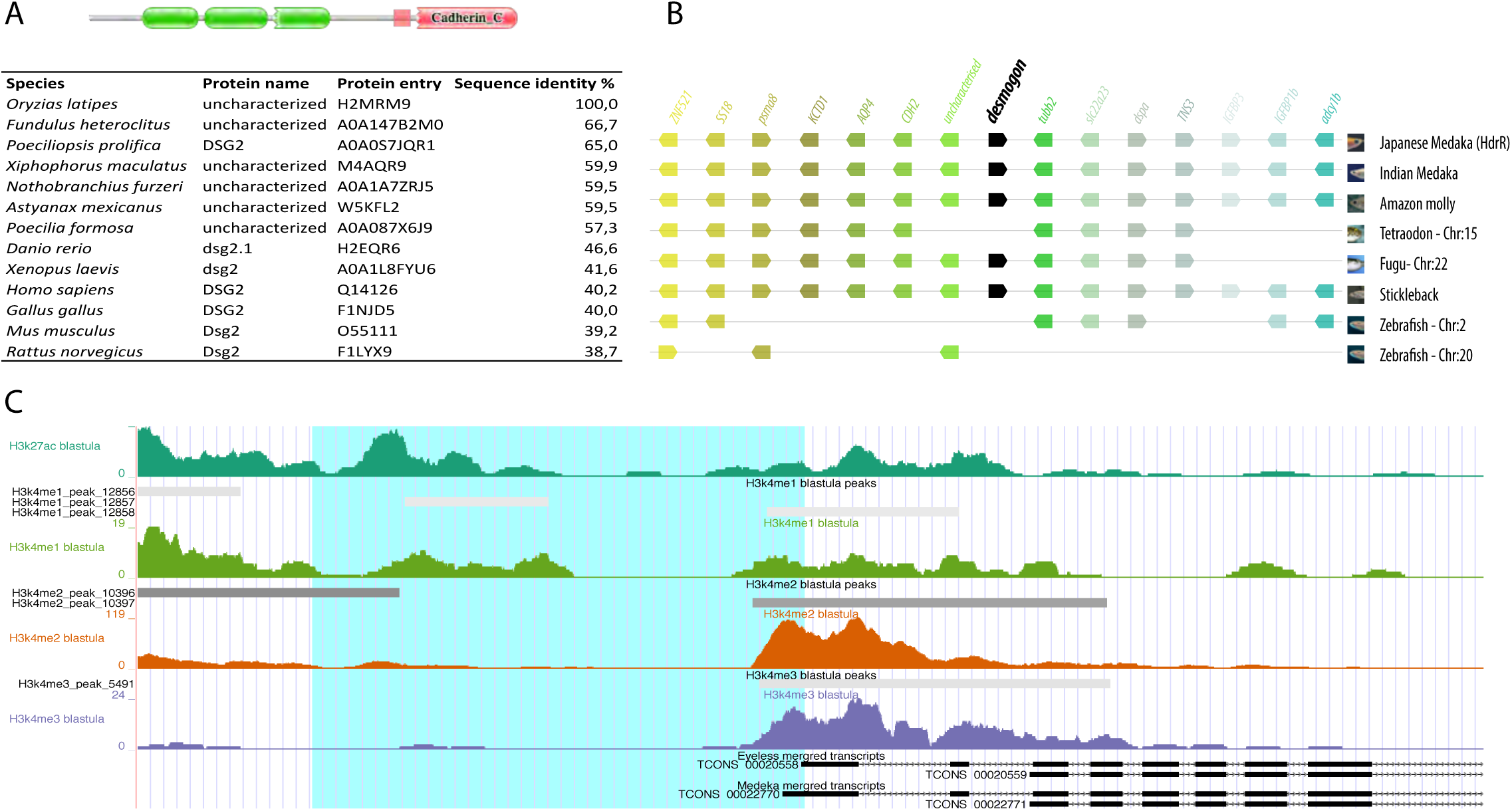
(A) Pfam predicted Desmogon protein domains. In green, 3 cadherin domains and in red, a cytoplasmic cadherin domain. Multiple sequence alignment of 12 selected species to uncharacterized Medaka protein(H2MRM9). Identity score reveals weak amino acid sequence conservation with closest hits. (B) Comparative genomic alignment of uncharacterized medaka transcript (ENSORLG00000017110) using GENOMICUS shows conservation of the *desmogon* locus in the vast majority of teleost branches; notice the loss of locus in Zebrafish and the Tetraodon. Locus of interest is highlighted by black *desmogon* label. Scheme modified from Genomicus to highlight syntenic genomic region. (C) Choosing of *desmogon* partial 2.2kb promoter, region highlighted in blue. H3K27ac, H3K4me1 and H3K4me2 peaks from UCSC genome browser Medaka blastula stage data at 2.2kb upstream of predicted *desmogon* TSS.

**Supplementary Figure 2.**
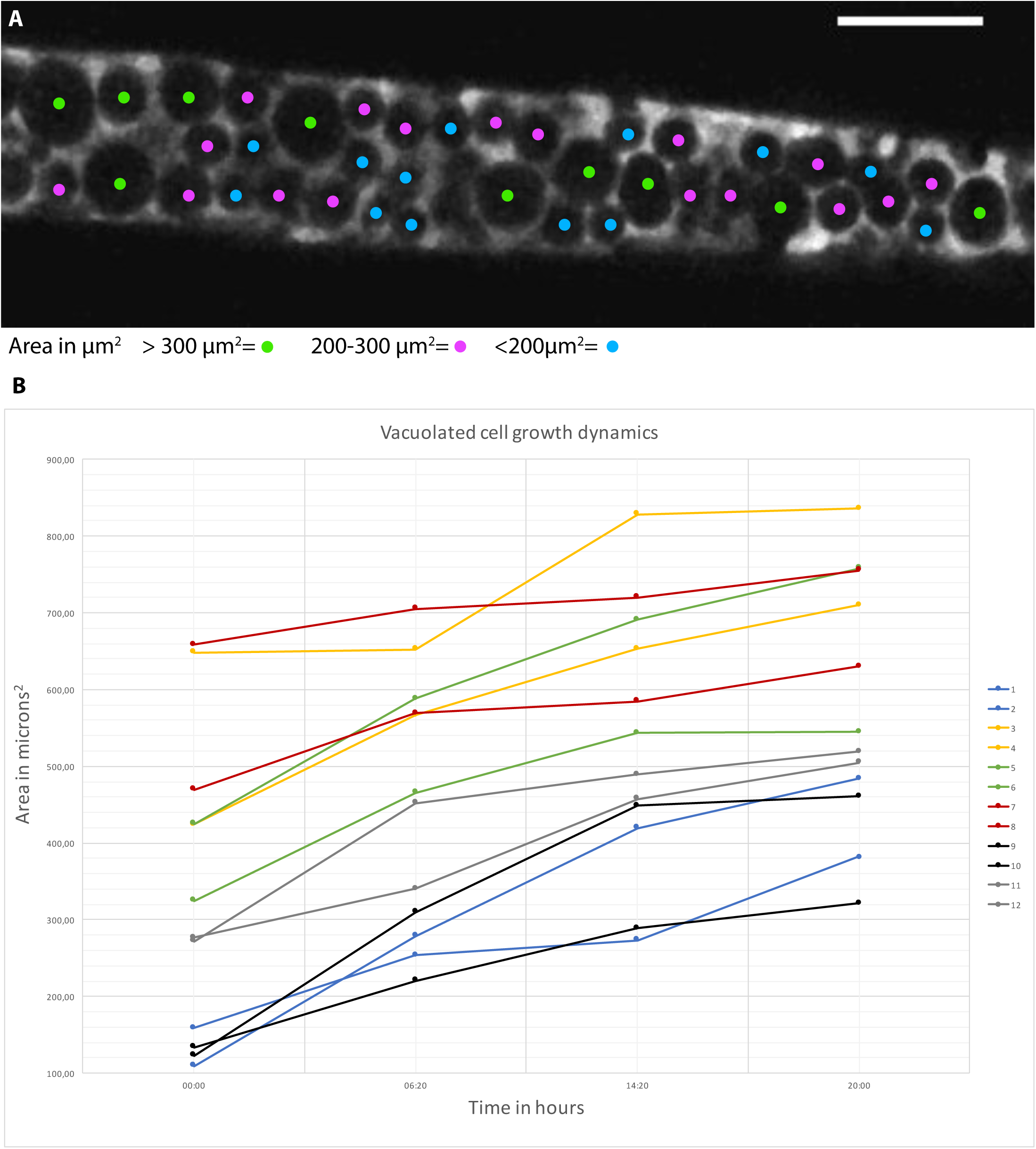
(A) Classification of vacuoles according to their area shows intermingled distribution of vacuolated cell size during notochordal growth in 3dpf Medaka notochords. (B) Area measurement on 12 paired vacuolated cells at 4 different time-point over a 20-hour period reveals the asynchronous nature of vacuolated cell growth. Neighbouring vacuolated cells share the same colour code and grow at different rates. Area was calculated on maximum projections using standard Fiji software.

**Supplementary Figure 3.**
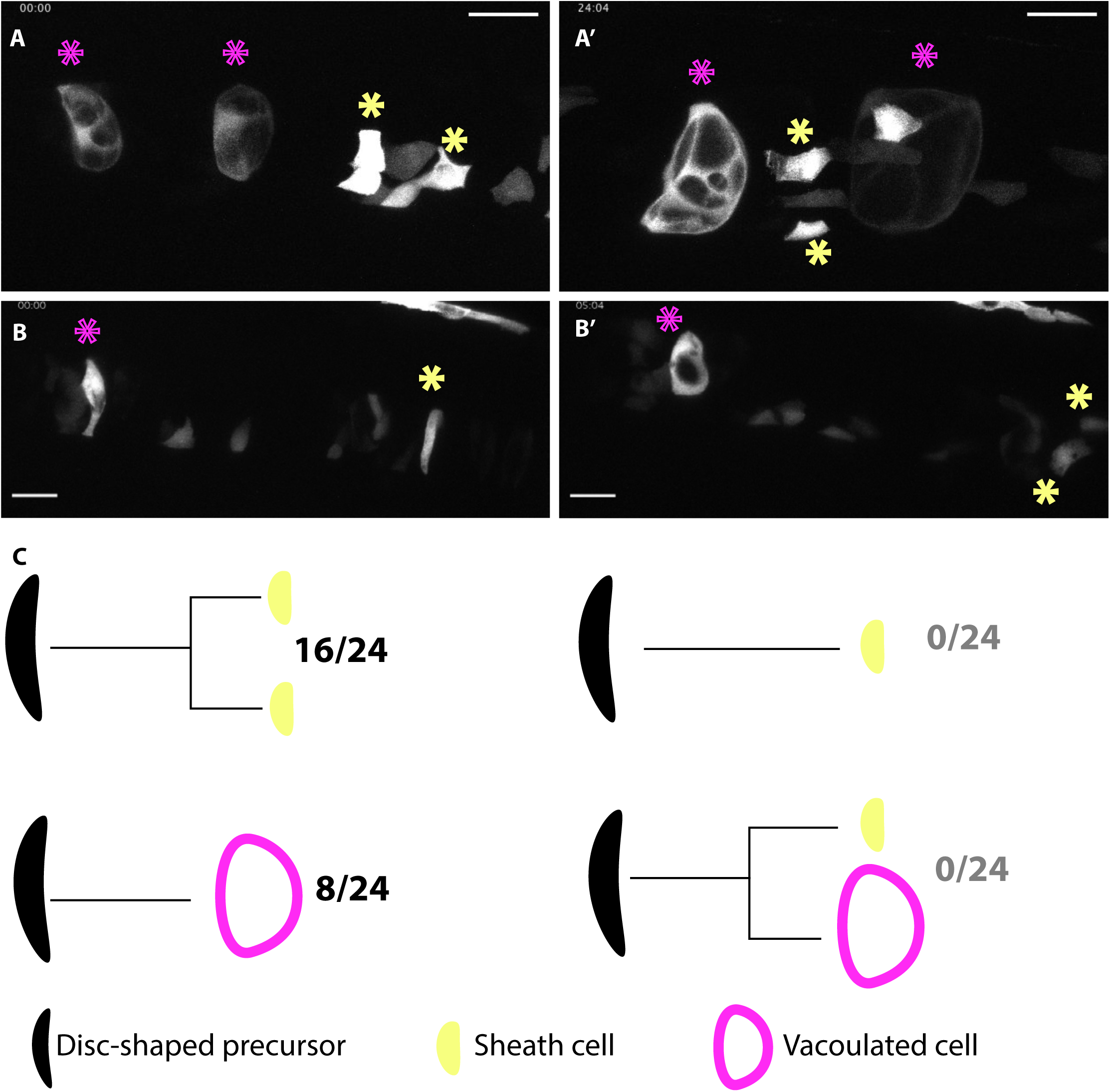
(A, A’) Time-lapse recording of clones of vacuolated cells (purple asterisks) and sheath cells (yellow asterisks) labelled after injection of *desmogon*:EGFP plasmid into Zebrafish embryos. Notice the growth of vacuolated cells over time is asynchronous. (B,B’) Time-lapse recording of notochord disc-shaped precursor cells labelled in injected zebrafish embryos. One labelled precursor directly trans-differentiates into vacuolated cell (purple asterisks) while the other undergoes a dorso-ventral division giving rise to two sheath cells (yellow asterisks). Scale bar= 30 microns. Time in hours. (C) Quantification of disc-shaped precursor behaviour during development of the notochord in zebrafish N= 4 embryos.

**Supplementary Figure 4.**
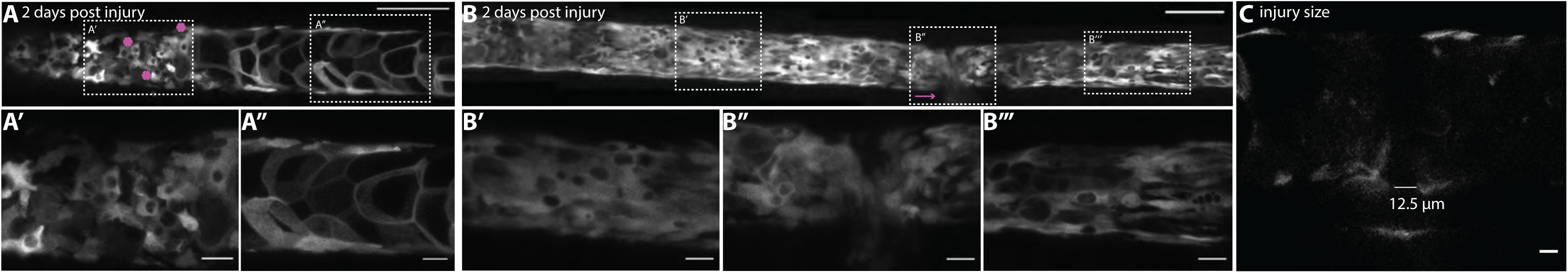
(A-A’’) Local injury response 2 days post injury. Locally restricted appearance of small vacuolated cells in the injured area. (B-B’’’) Global injury response 2 days post injury. Appearance of small vacuolated cells along the entire notochord. (C) Size of perinotochordal injury immediately after laser ablation.

**Supplementary Figure 5.**
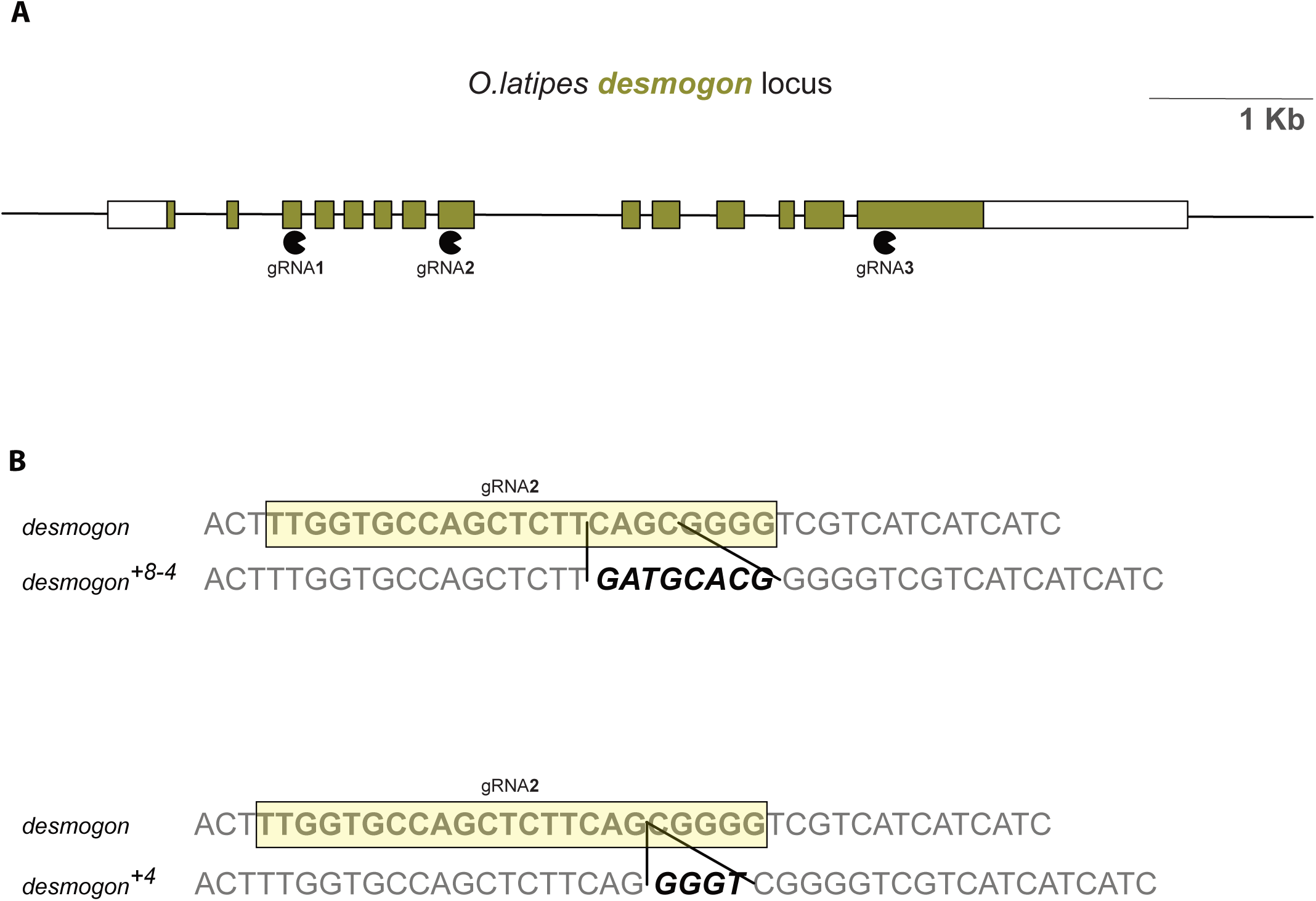
(A) Schematic view on the *desmogon* locus with exons targeted with gRNAs. (B) Mutant alleles with four base pair addition were isolated by sequencing and TIDE analysis.

**Supplementary Figure 6.**
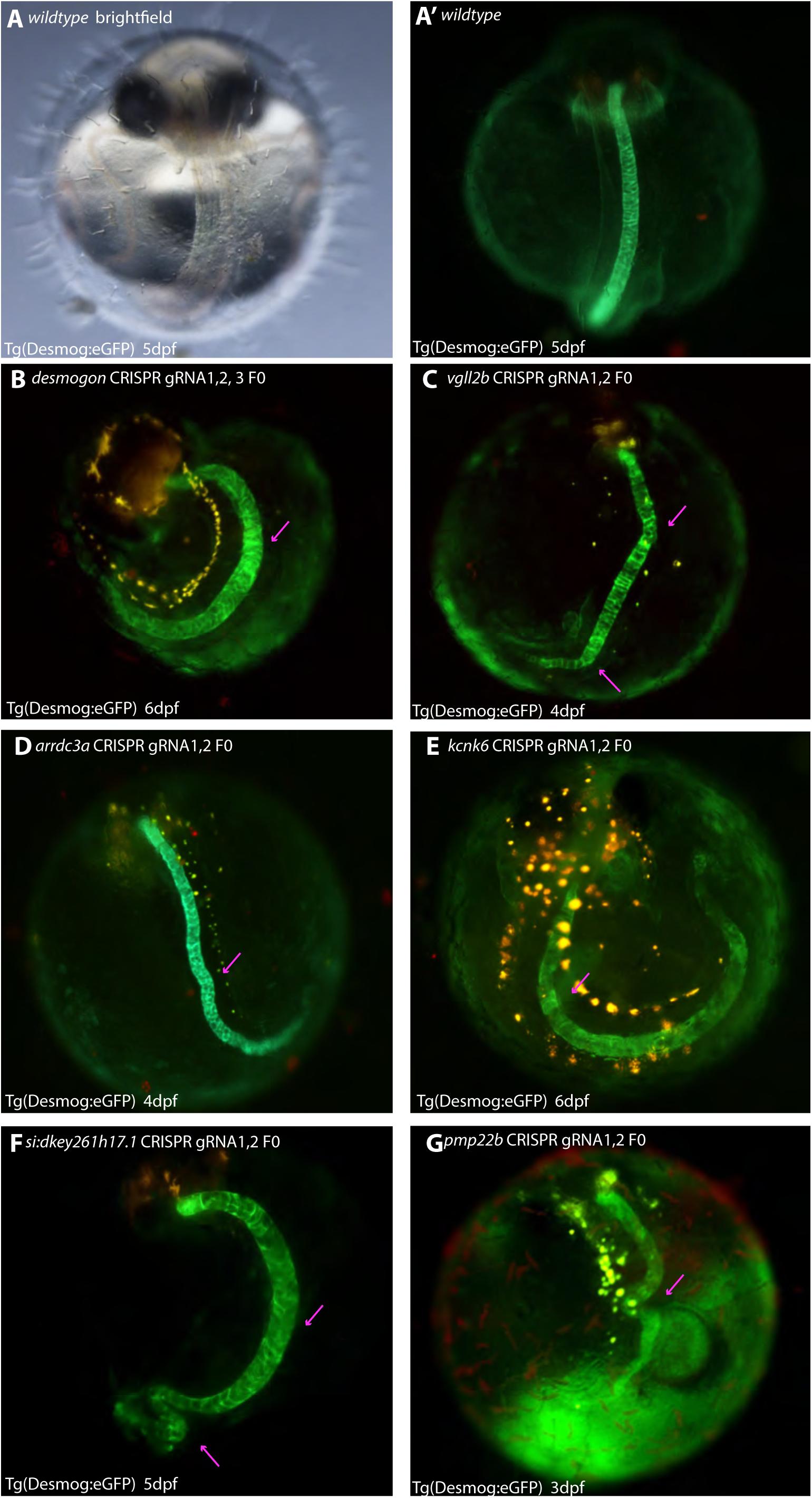
Embryonic phenotypes on F0 CRISPR screen. (A) Brightfield image of a wildtype medaka embryo. (A’) Tg(*desmog*:EGFP) wildtype embryo. (B) *desmogon* crispants show vacuolated cell lesions, magenta arrow highlights position of lesion. (C) *vgll2b* crispants show twisted and bent notochords, magenta arrows highlight position of major twists. (D*) arrdc3a* crispants show wavy and bent notochords (magenta arrows). (E) kcnk6 crispants show malformed and twisted notochords (magenta arrows). (F) *si:dkey261h17.1* show spirals and twisted notochords (magenta arrows). (G) *pmp22b* crispants show twisted and malformed notochords (magenta arrows).

## Materials and Methods

### Fish stocks and generation of transgenic lines

Medaka (*Oryzias latipes*) and Zebrafish (*Danio rerio*) stocks were kept in a fish facility built according to the local animal welfare standards (Tierschutzgesetz §11, Abs. 1, Nr. 1) as described before (Seleit et. al, 2017). Animal experiments were performed in accordance with European Union animal welfare guidelines.(Tierschutzgesetz 111, Abs. 1, Nr. 1, Haltungserlaubnis AZ35–9185.64 and AZ35–9185.64/BH KIT).

The strain used in this study is: Cab (medaka Southern wild type population). Tg(*desmogon*:EGFP) transgenic line was generated for this study by I-SceI mediated insertion, as previously described(Rembold et al. 2006). Briefly, a 2.2kb promoter region upstream of the desmogon CDS (but including the ATG of the first exon) was amplified from Medaka genomic DNA using forward primer (TCGCTGCTTGTTGTGTAGGT) and reverse primer (CATTGGCGCAGTGATTTGAA). The amplified fragment was subsequently cloned by A-tailing into a PGEMTeasy vector (Promega). From there it was sub-cloned into a vector with I-SceI sites and eGFP. This was then injected into Medaka embryos to obtain the transgenic line Tg(*desmogon*:eGFP).

### Imaging and Image analysis

Embryos were prepared for live-imaging as previously described (Seleit, et al. 2017a; Seleit, et al. 2017b) 20 mg/ml as a 20x stock solution of tricaine (Sigma-Aldrich, A5040-25G) was utilized to anaesthetise embryos. Glass-bottomed dishes (MatTek Corporation, Ashland, MA 01721, USA) contained the embryos that were in turn covered with low melting agarose (0,6% in ERM). Embryos were screened using an Olympus MVX10 binocular coupled to a Leica DFC500 camera. For image acquisition we used confocal microscopes Leica TCS SPE and Leica TCS SP5 II. Developing notochords were imaged for approximately 1 day using EMBL MuVi-SPIM (Krzic, Gunther, Saunders, Streichan & Hufnagel 2012b) with two illumination objectives (10x Nikon Plan Fluorite Objective, 0.30 NA) and two detection objectives (16X Nikon CFI LWD Plan Fluorite Objective, 0.80 NA). Embryos were placed in glass capillaries using 0,6% low melting agarose at room temperature. Standard Fiji software was used for all image analysis. Stitching was performed using 2D and 3D plugins on Fiji. For mosaic analysis, a *desmogon*:EGFP plasmid was injected embryos into 2 to 4 cell-stage embryos, which were grown until day 9 pf. For the quantification, we selected embryos bearing 1 and up to 4 clusters in the notochord, where clusters were defined as single cells or groups of cell that were at least one somite away from each other.

### Multi-photon Laser Ablations

A multi-photon laser was used in combination with a Leica TCS SP5 II microscope to perform ablations on vacuolated cells of the notochord and the peri-notochordal membrane in Tg(desmogon:eGFP). ‘Point ablations’ were chosen along the perimeter of each vacuolated cell. Laser of 880 nm wavelength was used and the power used ranged from 30-35% for 500ms (the time parameter was adjusted in the biological replicates when necessary). The targeted area was immediately imaged post-ablation to checked for signs of cellular bursting, debris and the release of GFP signal.

### CRISPR gRNAs and Cas9

Lab-made Cas9 mRNA was transcribed by mMachine Sp6 Transcription Kit (Thermo Fisher). The gRNAs for all targeted genes were designed in-silico using CCtop (Stemmer, Thumberger, et al. 2015). gRNA synthesis was done as previously reported (Stemmer, Thumberger, et al. 2015). All genes were targeted with 2 gRNAs against the CDS except *desmogon* where 3 gRNAs were designed and injected. The following is a list of gRNA sequences used in this study for each gene.

desmogon gRNA1 (CCCAUAUGGUGUGUUCAGCGUGG) gRNA2
(UUGGUGCCAGCUCUUCAGCGGGG) gRNA3 (AUCUAUGACUAUGAAGGUCGAGG)
vgll2b gRNA 1 (UGGGCCCCCAGACAUUCCUUCGG)
gRNA2(GGGUGCGCCCGUUUCACAGUGGG)
arrdc3a gRNA1(AAGUUCGUUGGACGGAAUCGAGG)
gRNA2(AUCCCGCAUGGUGGUCCCAAAGG)
kcnk6 gRNA1 (GUUCUCCAGCAUCGAGCGGCCGG) gRNA2
(UGUAUUGUAGUCGCCGGGCGAGG)
si:dkey-261h17.1 gRNA1 (GAGCUUGCUGUCACAAGCCUUGG)
gRNA2(CUGAGUCACUUCAAUCUGCCUGG)
pmp22b gRNA1(CCUGCUGCACAUUGCUGCACUGG) gRNA2
(CAGCUCUUCACCUUGCAGAAGGG)

Tg(*desmogon*:eGFP) were injected at the 1 cell stage with a solution containing 15 ng/μl of each gRNA for each gene individually and 150 ng/μl of CAS9 mRNA. The embryos were screened iteratively over the course of development for gross morphological phenotypes and for phenotypes in the notochord. For control injections two gRNAs targeting the *Oca2* locus and Cas9 (Lischik et al. 2018) were injected into Tg(*desmogon*:eGFP) embryos. In order to analyse the mutant alleles, genomic DNA was extracted from *wt* and *desmogon* mutant fish. A PCR flanking the gRNA2 locus was performed using primers *fwd* GCTGGCAGCCTTTGAAATTG *rev* TCGTACCTGACATTGGTGGC. The PCR products were sent to sanger sequencing and also cloned into TopoTA vector (Invitrogen) to distinguish single alleles. The analysis was complemented by using the online software tool TIDE (Brinkman et al., 2014), we were able to isolate the two alleles shown in Supplementary Figure 5.

### Whole-mount *in-situ* hybridization

A 604bp *in situ* probe for *desmogon* was generated from total cDNA of stage 33 medaka embryos by PCR using the following primers *fwd*: TTCTGCGAGATCAGGCTCAC *rev*: AAGGCCCCTCCTCTGTAACT and subsequently A-tailed and cloned into a PGEMTeasy vector (Promega). Sense and anti-sense probes were generated using Sp6 and T7 polymerases (Invitrogen) and the *In-situ* hybridization protocol was followed as previously reported in (Stemmer, Schuhmacher, et al. 2015). Hybridisations were performed overnight at 65°C then samples were incubated with an antibody against anti-digoxigenin conjugated with AP Fab fragments (1:2000; Roche, 11093274910). Staining was done using NBT/BCIP (Roche).

### Electron Microscopy sample preparation and imaging

10 dpf fish from Medaka wt Cab strain and stable Desmogon mutants were placed in a fixative consisting of 2.5% glutaraldehyde and 4% paraformaldehyde in 0.1M PHEM buffer. The fluorescing part of the notochord in mutant fish and equally small pieces of the wt fish was cut out in the fixative and fixation was continued for 30 min at room temperature and at 4°C overnight. The samples were further fixed in 1% osmium in 0.1M PHEM buffer, washed in water, and incubated in 1% uranylacetate in water overnight at 4°C. Dehydration was done in 10 min steps in an acetone gradient followed by stepwise Spurr resin infiltration at room temperature and polymerization at 60°C.The blocks were trimmed to get longitudinal sections of the notochord. 70nm thick sections were obtained using a leica UC6 ultramicrotome (Leica Microsystems, Vienna) and the sections were collected on Formvar-coated, copper slot grids and thereafter post-stained with uranyl acetate and Reynold‘s lead citrate. Imaging was done using a JEOL JEM-1400 electron microscope (JEOL, Tokyo) operating at 80 kV and equipped with a 4K TemCam F416 (Tietz Video and Image Processing Systems GmBH, Gautig).

### Transcriptomics data and candidate picking

The targeted genes were selected from single cell transcriptomics data generated by the Klein Lab (Briggs et al., 2018) based on a number of different criteria. First, high enrichment during early development of the notochord in zebrafish. Precisely, candidates were picked from 10, 14, 18 and 24 hpf data. Second, the selected genes were checked for conservation across vertebrates. Third the genes had to be well annotated but poorly characterized in both Medaka and Zebrafish. Lastly, a compact and short CDS, simple exon-intron structure/number and limited alternative splicing was favoured to improve chances of efficient mutagenesis with the CRISPR machinery.

### Bioinformatics tools

ENSEMBL(www.ensembl.org) was used to obtain all the coding sequences and upstream promoter regions for all genes analysed in this study. Supplementary Table 1 and 2 were generated using standard ensembl software using the 1-1 and 1-many orthologue comparisons. Genomic sequence alignments for synteny were performed on ensembl or using GENOMICUS (www.genomicus.biologie.ens.fr/). Protein information and amino acid sequence alignments were done using Uniprot (www.uniprot.org). UCSC (https://genome.ucsc.edu/) genome browser was used for the identification of the promoter region of desmogon. Pfam and PRINTS were used for information on protein domains and conserved motifs (https://pfam.xfam.org/) (http://130.88.97.239/PRINTS/index.php). Single cell transcriptomics data was visualized using (https://kleintools.hms.harvard.edu/paper_websites/wagner_zebrafish_timecourse2018/springViewer.html?coarse_grained_tree) and the selected candidates were obtained from list of enriched genes in developing Zebrafish notochords.

## Acknowledgments

We would like to thank Sylvia Urbansky and Steffen Lemke for critical input on the manuscript. J. Wittbrodt for the use of confocal and light sheet microscopes and generous support. We would like to thank C. Funaya, S. Gold and S. Hillmer from the EMCF facility at Heidelberg University for the preparation, imaging and processing of the electron microscopy data. We would like to thank Nihad Softic for injections for spare labeling, Pui Oui Hong for help with the genotyping of *desmogon* mutants, and Carina Vibe for the medaka bone staining protocol. We thank E. Leist, M. Majewski and A. Saraceno for fish husbandry.

## Funding

### Deutsche Forschungsgemeinschaft (SFB 873)

- Ali Seleit
- Karen Gross
- Lazaro Centanin

The funders had no role in study design, data collection and interpretation, or the decision to submit the work for publication.

